# swCAM: estimation of subtype-specific expressions in individual samples with unsupervised sample-wise deconvolution

**DOI:** 10.1101/2021.01.04.425315

**Authors:** Lulu Chen, Chiung-Ting Wu, Chia-Hsiang Lin, Rujia Dai, Chunyu Liu, Robert Clarke, Guoqiang Yu, Jennifer E. Van Eyk, David M. Herrington, Yue Wang

## Abstract

**Motivation:** Complex biological tissues are often a heterogeneous mixture of several molecularly distinct cell or tissue subtypes. Both subtype compositions and expressions in individual samples can vary across different biological states or conditions. Computational deconvolution aims to dissect patterns of bulk gene expression data into subtype compositions and subtype-specific expressions. Typically, existing deconvolution methods can only estimate averaged subtype-specific expressions in a population, while detecting differential expressions or co-expression networks in particular subtypes requires unique subtype expression estimates in individual samples. Different from population-level deconvolution, however, individual-level deconvolution is mathematically an underdetermined problem because there are more variables than observations.

**Results:** We report a sample-wise Convex Analysis of Mixtures (swCAM) method that can estimate subtype proportions and subtype-specific expressions in individual samples from bulk tissue transcriptomes. We extend our previous CAM framework to include a new term accounting for between-sample variations and formulate swCAM as a nuclear-norm and ***ℓ*_2,1_**-norm regularized matrix factorization problem. We determine hyperparameter values using a cross-validation scheme with random entry exclusion and obtain a swCAM solution using an efficient alternating direction method of multipliers. The swCAM is implemented in open-source R scripts. Experimental results on realistic simulation data show that swCAM can accurately estimate subtype-specific expressions in individual samples and successfully extract co-expression networks in particular subtypes that are otherwise unobtainable using bulk expression data. Application of swCAM to bulk-tissue data of 320 samples from bipolar disorder patients and controls identified changes in cell proportions, expression and coexpression modules in patient neurons. Mitochondria related genes showed significant changes suggesting an important role of energy dysregulation in bipolar disorder.

**Availability and implementation:** The R Scripts of swCAM is freely available at https://github.com/Lululuella/swCAM. A user’s guide and a vignette are provided.

**Supplementary information:** Supplementary data are available at *Bioinformatics* online.

## 1 Introduction

Alteration of biological processes in particular cell or tissue subtypes may lead to development of disease (Fan, et al., 2020; Herrington, et al., 2018). In order to understand molecular mechanisms affecting disease it is critical to study molecularly distinct cell or tissue subtype-specific effects in the context of a complex ecosystem (Chasman and Roy, 2017). However, these specific mechanisms are typically not revealed when a heterogeneous tissue is studied in bulk (Parker, et al., 2020). Computational deconvolution is a data-driven cost-effective technique to dissect patterns of bulk-tissue expression data into subtype compositions and subtype-specific expressions (Avila Cobos, et al., 2018; Chen, et al., 2020; Wang, et al., 2016). Importantly, unsupervised deconvolution of bulk tissue expression profiles has the advantage of observing subtype-specific expression patterns in the context of cell-cell and cell-matrix interactions that define the local tissue environment compared with single-cell or cell-sorted data where the effects of these interaction may be lost (Herrington, et al., 2018). Moreover, computational deconvolution has the potential of extracting novel yet reproducible biological information from vast amounts of existing bulk transcriptome data, resulting in gains in statistical power and opportunities for further insights for a wide range of previous experimental designs (Wang, et al., 2016).

Typically, existing deconvolution methods can only estimate averaged subtype expressions that are shared in a population (Avila Cobos, et al., 2018; Wang, et al., 2016), while detecting differential expressions or co-expression networks in particular subtypes requires subtype-specific expressions in individual samples (unique for each individual) (Zhang and Horvath, 2005; Zhang, et al., 2009). When multiple measures of bulk-tissue expression from the same individuals are available, population-level deconvolution methods, such as Convex Analysis of Mixtures (CAM - unsupervised) (Wang, et al., 2016) or Multimeasure Individual Deconvolution (MIND - supervised) (Wang, et al., 2020), can be readily applied but with reduced statistical power and subtype-resolution. Correspondingly, some semi-supervised methods have recently been proposed to exploit single-measure bulk data, including Tensor Composition Analysis (TCA) on DNA methylation data (Rahmani, et al., 2019), CIBERSORTx and Bayesian MIND (bMIND) on gene expression data (Newman, et al., 2019; Wang, et al., 2020). TCA works specifically on DNA methylation data, based on an assumed model similar to MIND, and requires *a priori* knowledge or estimate of subtype proportions, CIBERSORTx relies on subtype expression signatures derived from single-cell or bulk-sorted reference profiles and uses pseudo non-negative least squares to achieve high-resolution expression purifications leveraging grouped sample structures, and bMIND uses again information from scRNA-seq data fully, as prior information, to refine subtype expression estimates per bulk sample.

To address the critical problem of the absence of fully unsupervised individual-level deconvolution methods, we develop a sample-wise Convex Analysis of Mixtures (swCAM) method to estimate constituent proportions and subtype-specific expressions in individual samples from bulk-tissue data (unique for each individual). We extend our previous CAM framework to include a new term that accounts for subtype variation among samples. Because individual-level deconvolution is mathematically underdetermined with more variables than observations, we formulate swCAM as a nuclear-norm and *ℓ*_2,1_-norm regularized low-rank matrix factorization problem (Hastie, et al., 2015). We obtain the swCAM solution using a modified and efficient alternating direction method of multipliers (ADMM) in which hyperparameter values are determined by a cross-validation scheme with random entry exclusion (Chi, et al., 2017). Specifically, the nuclear-norm regularization term focuses on modeling the between-sample subtype variation of low-rank gene co-expression function modules (Zhang and Horvath, 2005), and encourages a unique and biologically plausible solution (Supplementary Information). Experimental results on realistic simulation data sets show that swCAM can accurately estimate subtype-specific expressions of major subtypes in individual samples and successfully extract co-expression networks in particular subtypes that are otherwise unobtainable using bulk expression data. The swCAM tool enables statistically-principled large-scale subtype-level downstream analyses, such as detecting differentially expressed genes or differential dependency networks in particular subtypes.

## 2 Method

### 2.1 Problem formulation of swCAM

Population-level unsupervised deconvolution by CAM considers a classical linear latent variable model 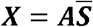, where ***X*** is the bulk-level expression matrix (sample vs gene), ***A*** is the sample-wise subtype proportion matrix (sample vs subtype), and 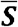 is the population-level subtype expression matrix shared across individuals (subtype vs gene) (Chen, et al., 2020; Wang, et al., 2016). To achieve individual-level unsupervised deconvolution, we extend the existing model to include a new term ***T*** that accounts for subtype variation among samples, see Figure 1. Specifically, we define the sample-wise subtype expression matrix as 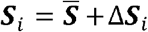, where Δ***S**_i_* is the subtype variation matrix associated with sample *i*. Correspondingly, the bulk-level expression ***x**_i_* associated with sample *i* is given by

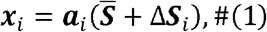

where ***α**_i_*, is the *i*th row vector of subtype proportion matrix.

**Figure 1.**
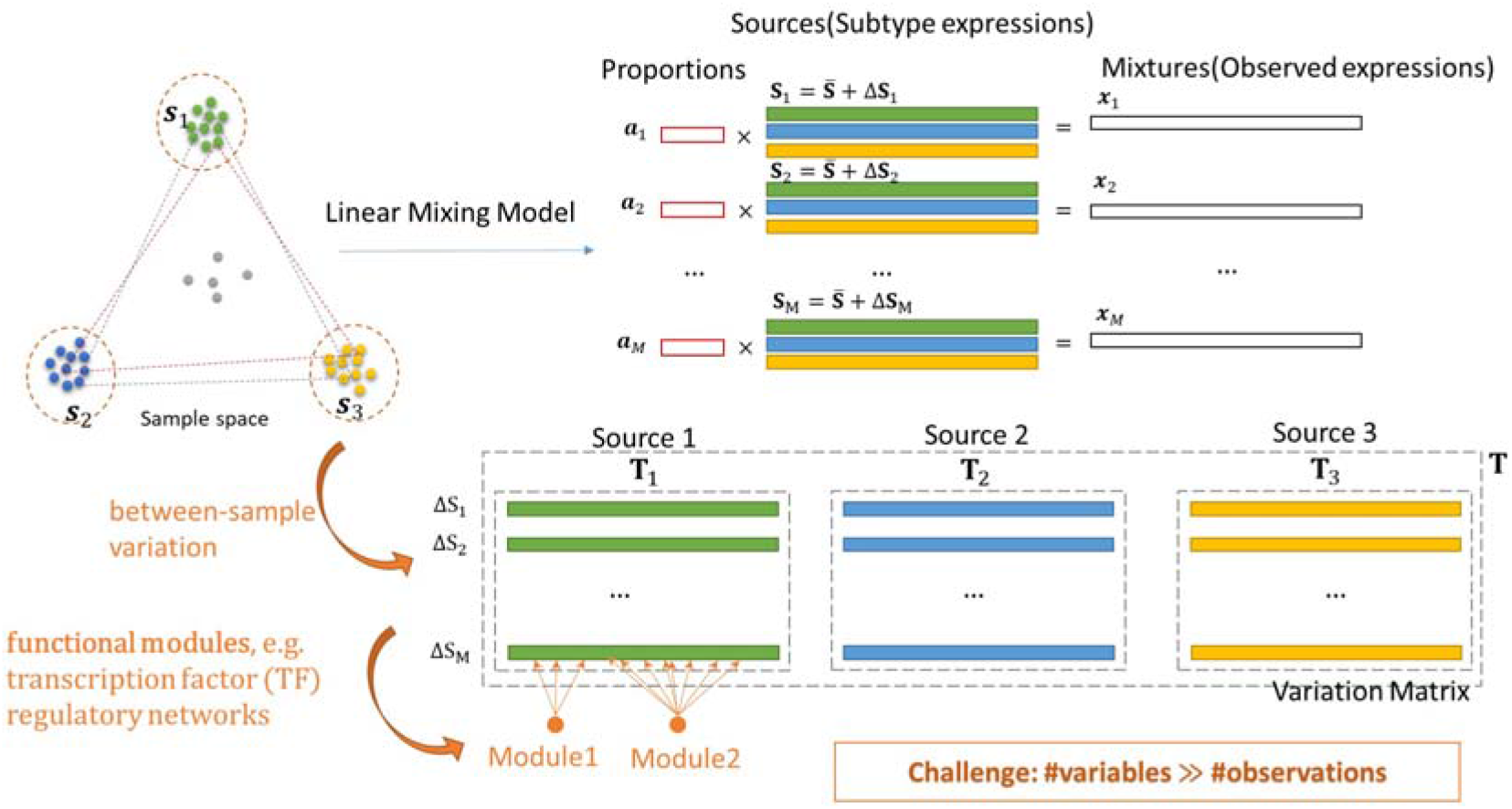
Illustrative diagram of swCAM framework (***T*** matrix is the transposed version of its text notation for simplicity sake).

Let 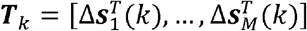 host the fcth columns in all Δ***S**_i_* associated with subtype *k*, and assume ***T**_k_* is a low-rank structured sparse matrix (Hastie, et al., 2015), the swCAM approach seeks to simultaneously estimate the subtype variation matrix Δ**S**_*i*_ for all samples by optimizing the following objective function, given the estimated ***A*** and 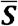 by CAM (Chen, et al., 2020; Wang, et al., 2016):

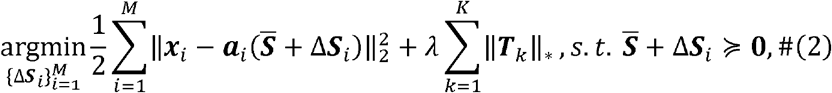

where *M* is the number of samples, *K* is the number of subtypes, ║ ║_2_ denotes L_2_ norm, ║ ║, denotes nuclear norm, and *λ* is the regularization hyperparameter. The premise behind a nuclear norm regularized subtype variation term ***T**_k_* is that there is only a small number of molecular function modules influencing the subtype variation and other noise-nature variations are of no biological meaning (Zhang and Horvath, 2005).

### 2.2 Optimization of swCAM objective function

The swCAM objective function given in (2) is convex with respect to the block-wise variables 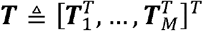 or 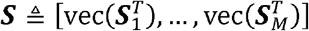, where vec(.) is the matrix vectorization operator (Supplementary Information). We propose to solve (2) by adapting the computationally efficient ADMM strategy and its default mathematical notations (Chi, et al., 2017), naturally decoupling the non-smooth regularization term from the smooth loss term. Specifically, we reformulate (2) into a new form where the primal variable is “split” into several parts, with the associated objective function “separable” across such splitting (Chi, et al., 2017).

Let 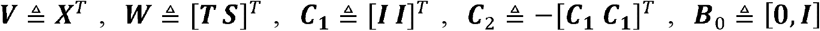, 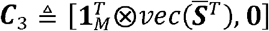 (⊗ denotes the Kronecker product), and 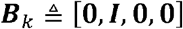, with *k* = 0…., *K*. We can simplify (2) into its equivalent form, given below

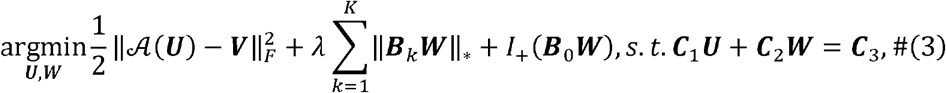

where *I*_+_(·) is the indicator function for the non-negative orthant; *I*_+_(***B***_0_***W***) = *I*_+_(***S***) = 0 if ***S*** ≽ **0**_*KL*×*M*_ (*I*_+_(*U*) ^=^ +∞, otherwise), see Figure 2. The linear transformation 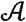 in the first term is 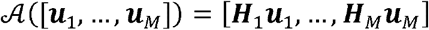 with ***H**_i_* = [***α**_i_*⊗*I_L_*], *i* = 1,…, *M*. Note that the two split block variables ***U*** and ***W*** in (3) is associated with the ADMM framework and notations (Boyd, et al., 2011). When (3) is solved, the solution ***T*** of (2) can be readily extracted from ***W*** (Supplementary Information).

**Figure 2.**
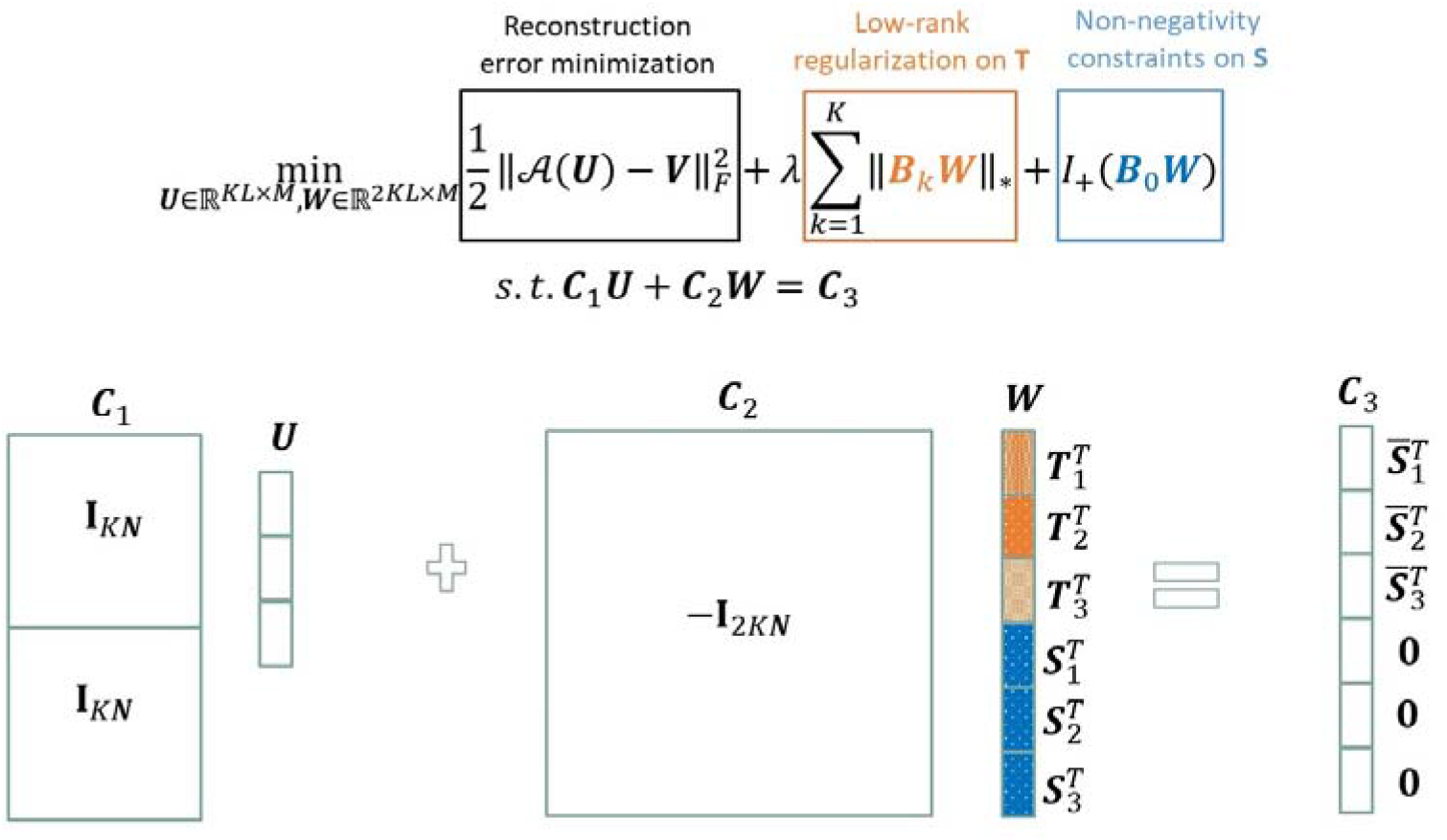
The objective function of swCAM for sample-specific deconvolution problem and its reformulation by ADMM. (For convenient illustration, *T* matrix in all figures are the transposed version of those in the text and equations.)

### 2.3 Regularization of between-sample variation

Nuclear-norm based regulation on between-sample variation ***T*** aims to detect unique and meaningful patterns of interest for each individual sample in reference to noise effect. When setting hyperparameter *λ*, a larger value will coerce overall between-sample variation to be zero, while a lower value will encourage stronger subtype-specific between-sample variation.

Cross-validation is a popular and effective strategy for tuning hyperparameters to reach a good balance between underfit and overfit. In a classical setting, one round of crossvalidation excludes a certain portion of samples and uses the model learned from the remaining samples to predict the excluded ones. Then every model is assessed by summarizing prediction performances across multiple rounds. In swCAM, however, the estimate of subtype expression in individual samples cannot be directly used to predict the excluded samples. Alternatively, we propose to randomly exclude entries rather than samples in ***X*** matrix, see Figure 3, a strategy similar to the one widely used in missing value imputation. Specifically, we randomly remove some entries in ***X*** matrix and accordingly rewrite the objective function (2) as

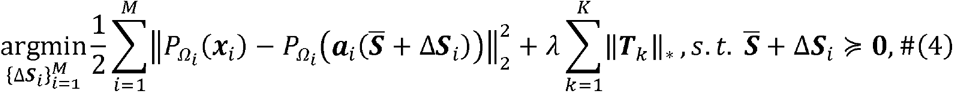

where *P*_Ω_*i*__(*x_i_*) denotes the modified ***x**_i_* vector with its entries in Ω_*i*_ untouched while others set to zero. The workflow of 10-fold cross-validation is summarize as follows:

1. Randomly split all entries into 10 folds of roughly equal size;
2. Remove onefold and use the remaining 9-folds to solve (4) with varying values of *λ* ∈ [*λ*_1_, *λ*_2_,…];
3. Use estimated Δ***S**_i_*(*λ*) to reconstruct ***X*** and record only the predicted values;
4. Repeat (2)-(3) steps to reconstruct 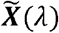 with all missing entries predicted;
5. Calculate root-mean-square error (RMSE)

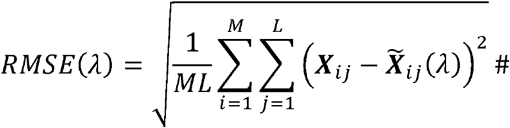

**Figure 3.**
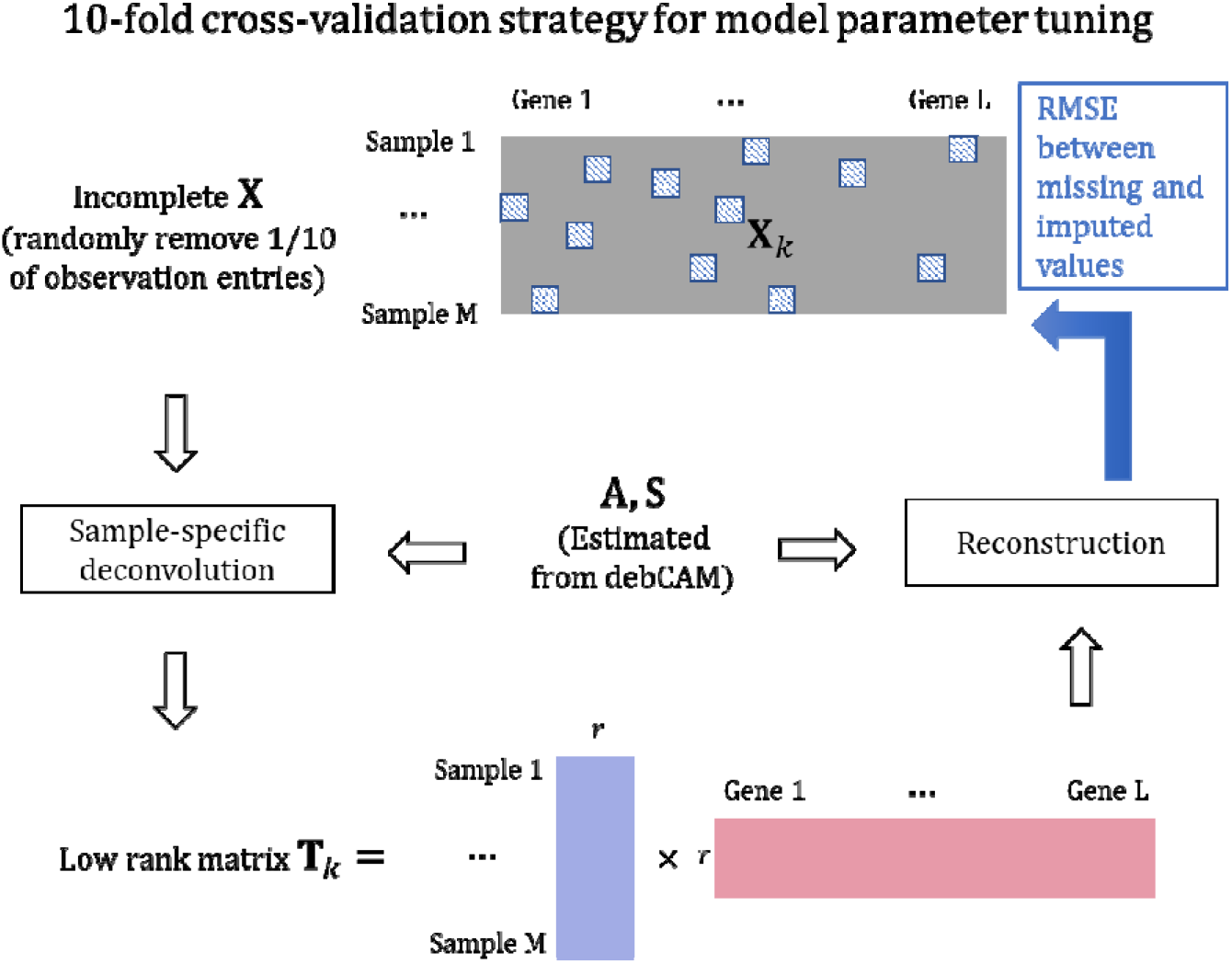
10-fold cross-validation strategy for model parameter tuning. A part of entries is randomly removed before applying swCAM. The removed entries are reconstructed by estimated matrix and compared to observed expressions for computing RMSE to decide the optimal hyperparameter.

Determine the optimal value *λ*_*_ yielding the minimum RMSE. Warm start can be used in step (2) with a decreasing sequence of λ. Again, the optimization problem (4) can be solved by the ADMM algorithm (Supplementary Information).

### 2.4 Sparsity regularization

Based on the further assumption that only a small number of genes are involved in each of functional modules, we propose to impose a row-sparsity regularization using *ℓ*_2,1_-norm on the between-sample variation term ***T***. The alternative swCAM objective function becomes, *s.t*. 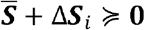,

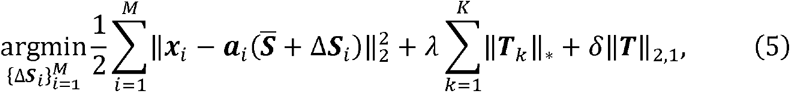

where *δ* > 0 is the regularization hyperparameter. The optimization problem (5) can be solved by the extended ADMM algorithm (Supplementary Information).

## 3 Results

We first conducted two phased computational experiments to assess the performance of swCAM method and R script for fully unsupervised sample-wise deconvolution, namely validation on ideal simulation and assessment on realistic simulation. The workflow of swCAM is implemented as an open-source R scripts together with the existing debCAM package (Chen, et al., 2020). The simulation datasets are generated from a benchmark real gene expression dataset (GSE19380) (Kuhn, et al., 2011). The ideal simulation assumes an independent relationship between variance and mean for genes, and the realistic simulation entertains a variance-mean relationship for genes close to that observed in real gene expression data. We then conducted biomedical case studies to demonstrate the real-life utility of swCAM tool in biomedical research.

### 3.1 Validation on ideal simulation

The ideal simulation involves three subtypes, twelve function modules (four unique functional modules in each subtype), 300 genes, and 50 samples. The baseline subtypespecific expression profiles 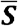 are sampled from the real gene expression of the purified subtypes in GSE19380, the standardized sample-specific proportion vectors ***a_i_*** are drawn randomly from a flat Dirichlet distribution, and the sparse between-sample variation matrix Δ***S**_i_*(*k,f*) is formed by assigning a value to *j*th gene from normal distribution 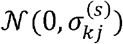 only if it participates in an unique function module in *k*th subtype (Figure 4). Each functional module is formed a by highly correlated gene pairs (co-expression network), 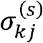 are drawn from the uniform distribution u[50,300], and the experimental noise ***n**_i_* is drawn from the zero-mean normal distribution with 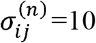. We tested swCAM on a large number of simulation datasets with varying hyperparameter settings.

**Figure 4.**
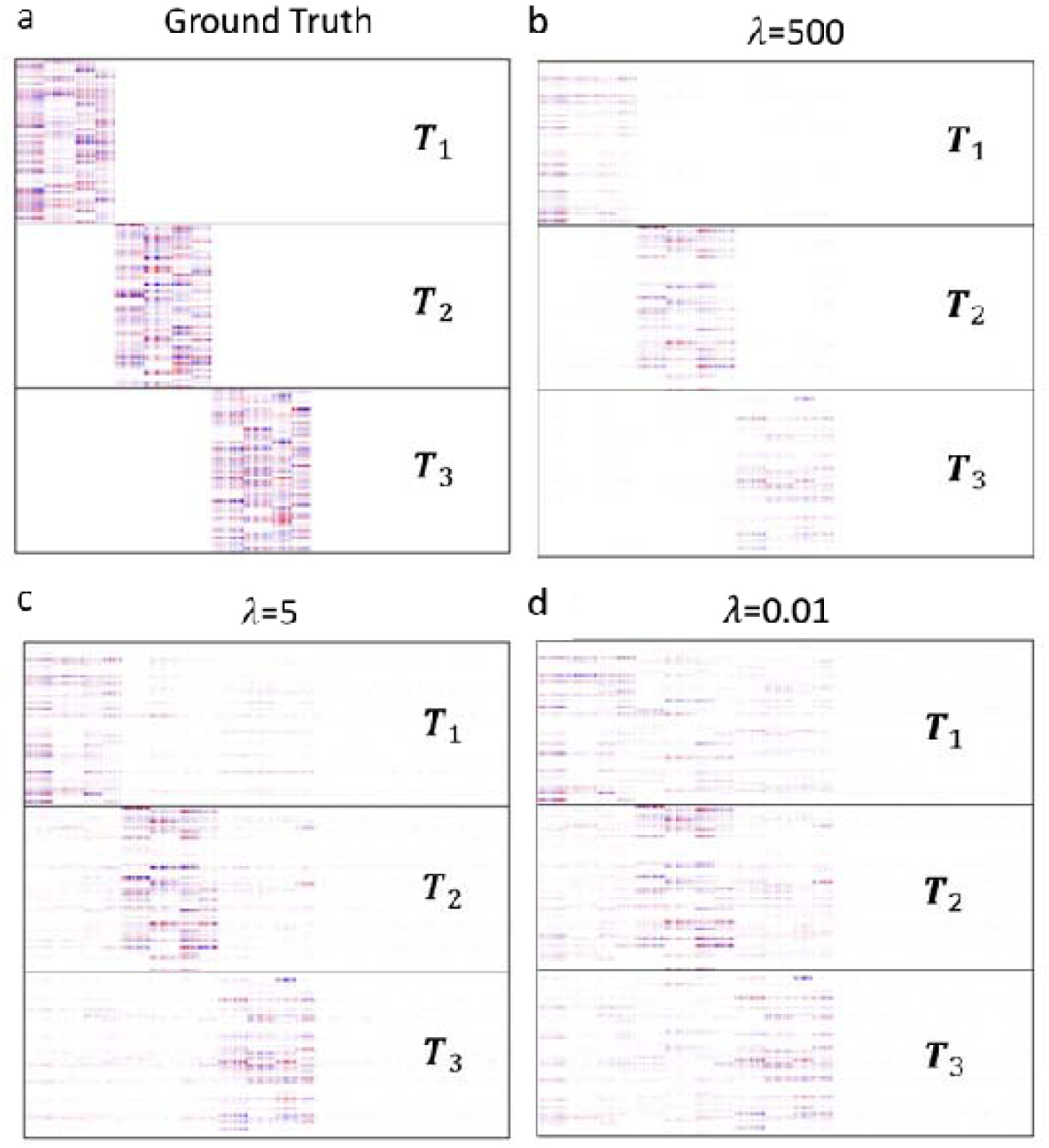
Heatmap of matrix estimate compared to ground truth in ideal simulation with varying.

The experimental results show that swCAM can blindly and successfully detected all twelve structured between-sample variation patterns when the regularization hyperparameter value *λ* is properly set by the proposed cross-validation guideline, see Figure 4 with given ground truth. Specifically, the RMSE obtained by 10-fold cross-validation is relatively small when *λ* = 1~50 and reaches the minimum at *λ* = 5. This event unsurprisingly coincided with the point where both primal and dual residuals diminish in the ADMM algorithm, see Figure 5 (Supplementary Information). Moreover, our experiments show that swCAM can extract more accurate between-sample variations from high abundant subtypes as compared with low abundant subtypes, see Figure 5 (green points - low abundant subtypes, red points – high abundant subtypes), where the Pearson’s correlation coefficients of the estimate and ground truth are 0.6079, 0.8329, 0.9412, and 0.9550, corresponding to the four quartiles of abundance level, respectively. This observation is consistent with what reported in the literature (Rahmani, et al., 2019; Wang, et al., 2020).

**Figure 5.**
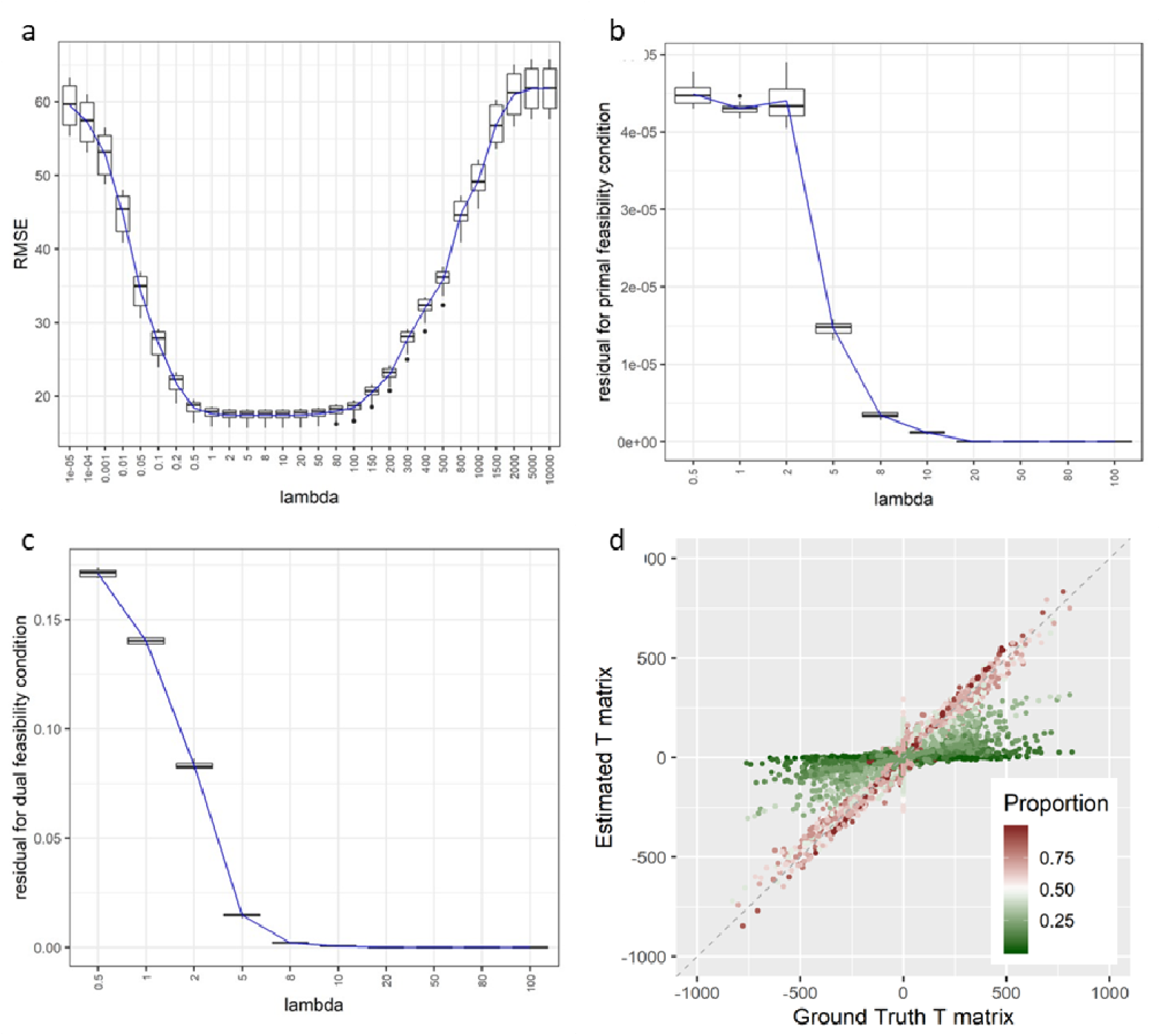
10-fold cross-validation results with different values in the ideal simulation. (a) RMSE curve; (b) Residuals for primal feasibility condition; (c) Residuals for dual feasibility condition; (d) Color-coded correlation scatter plot on the matrix entries between swCAM estimate and ground truth influenced by the abundant level of subtypes in the samples.

### 3.2 Assessment on realistic simulation

We modified the same design of ideal simulation by incorporating the variance-mean relationship widely observed in real gene expression data. First, the variance 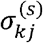 of subtype expressions and variance 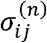 of overall noise are set to be proportional to the means of subtype and mixed expressions, respectively. Second, the coefficient of variation (ratio of standard deviation to mean) is drawn from the uniform distributions u[0.15,0.3] and u[0.02,0.05], respectively.

The experimental results show consistent trends and comparable performance of swCAM, clearly revealing all twelve structured between-sample variation patterns, see Figure 6. However, due to the impact of varying noise levels, the between-sample variation patterns estimated by swCAM expectedly becomes blurred and noisier. Our experimental results also show that the additional *ℓ*_2,1_-norm based sparsity regularization can jointly produce a much cleaner estimate of the between-sample variation patterns. Again, Figure 6d shows that the between-sample variations can be estimated more accurately from high abundant subtypes as compared with low abundant subtypes, where the Pearson’s correlation coefficients of the estimate and ground truth are 0.1979, 0.2223, 0.3683, and 0.7074, corresponding to the four quartiles of abundance level, respectively (Supplementary Information).

**Figure 6.**
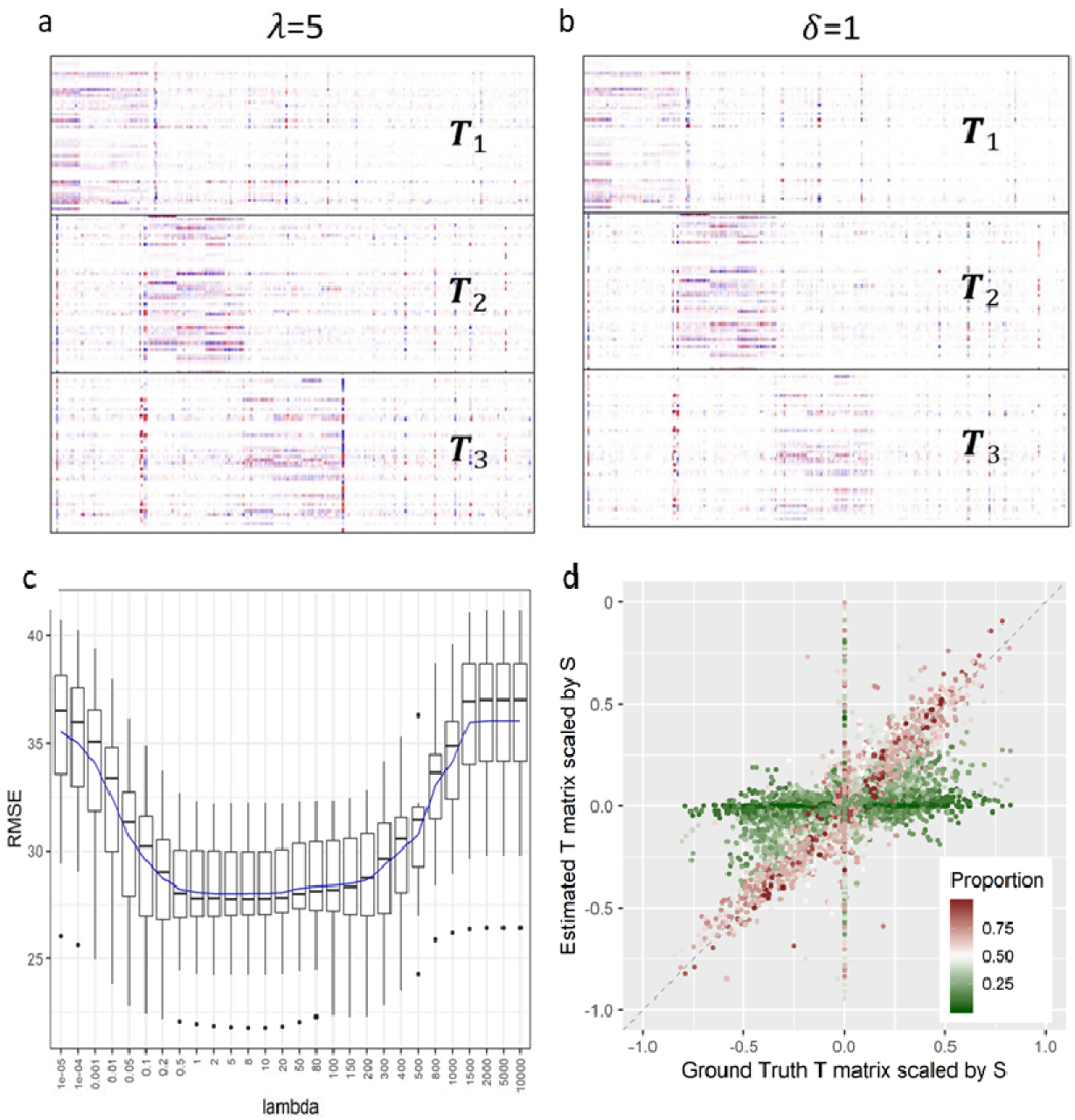
Experimental results obtained from the realistic simulation. (a-b) Heatmaps of matrix estimate with proper values of and. (c) 10-fold cross-validation RMSE curve. (d) Color-coded correlation scatter plot on the matrix entries between swCAM estimate and ground truth influenced by the abundant level of subtypes in the samples.

To further test the utility of swCAM in subsequent network analysis targeting the twelve functional modules, we performed the Weighted Gene Correlation Network Analysis (WGCNA) (Langfelder and Horvath, 2008; Zhang and Horvath, 2005) using the sample-wise subtype-specific expression profiles estimated by swCAM. The experimental results show that while the estimated between-sample variation patterns were less ideal as compared to the ground truth, the results of swCAM (as the input to WGCNA) enabled WGCNA to successfully reconstruct the weighted gene correlation networks associated with the twelve functional modules, see Figure 7 (Supplementary Information). Specifically, swCAM-WGCNA accurately identified the exact four true gene co-expression modules with very few missing genes resided within the second and third subtypes, and minor false positives in the first subtype. In contrast, without swCAM based deconvolution, WGCNA using the bulk expression data cannot detect any of the embedded functional modules, but detected three false modules reflecting the confounding effect of the varying subtype proportions.

**Figure 7.**
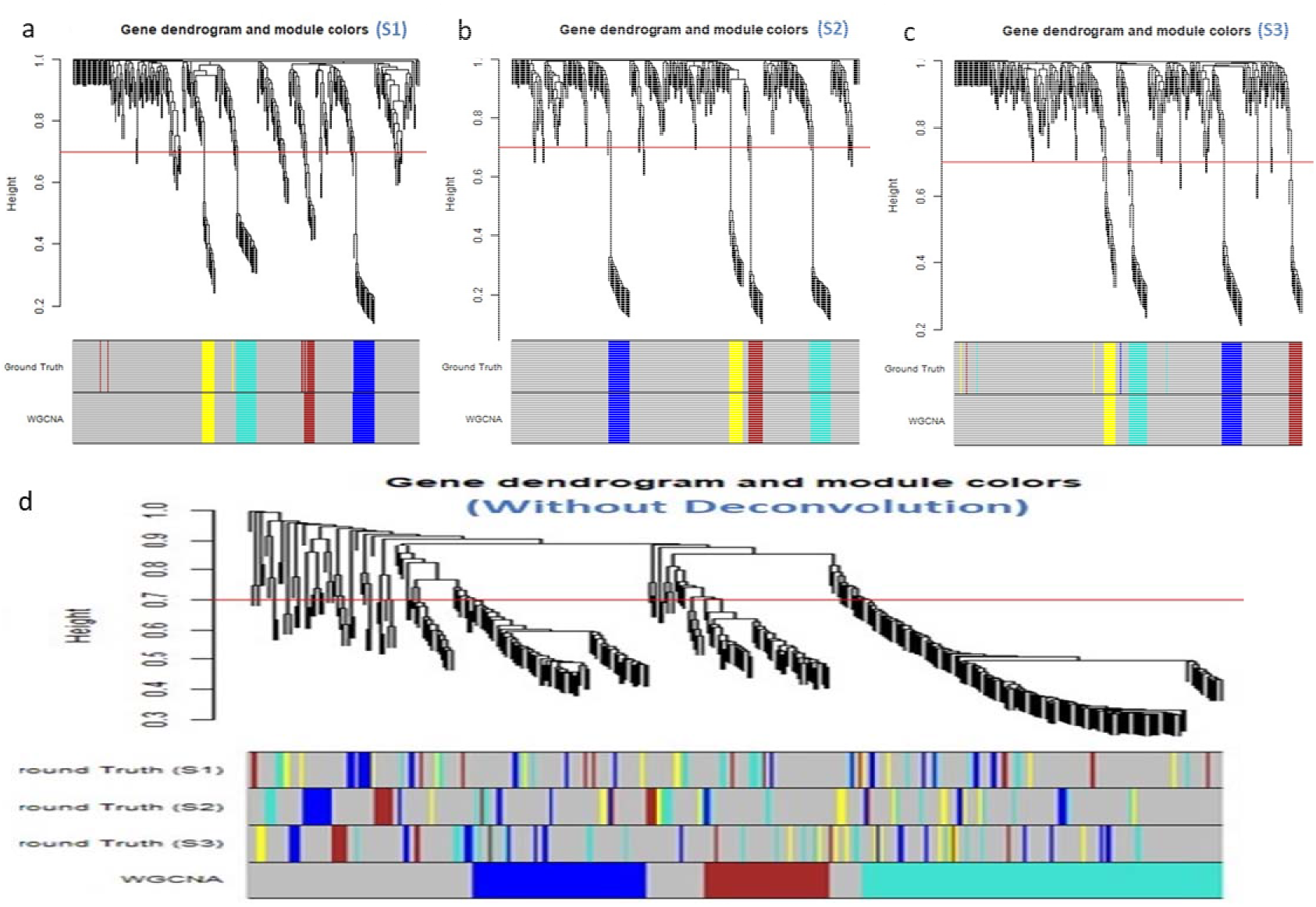
swCAM-WGCNA analysis. (the network interconnectedness is measured by topological overlap; cutHeight = 0.7; minSize = 8). (a-c) Gene co-expression modules detected by WGCNA using swCAM estimated sample-specific expression profiles in individual subtypes, with λ=5 and δ=1. (d) Gene co-expressed function modules detected falsely by WGCNA using bulk gene expression data.

### 3.3 Real-life biomedical case study

To further demonstrate biomedical utility of the swCAM method, we applied the swCAM software tool to enable detecting subtype-specific differential expressed genes and coexpression networks from RNA-seq data of human brains. The analytic pipeline is given in Figure 8a. This experimentally-acquired RNA-seq dataset was obtained from a cohort of “controls” (n=248) and “bipolar disorder (BD)” (n=72) human brain specimens located at dorsolateral prefrontal cortex (DLPFC) (Gandal, et al., 2018). The gene expression levels were quantified in Fragments Per Kilobase of transcript per Million mapped reads (FPKM). After quality control, 14,865 genes were retained, and expression profiles were normalized. Potential confounders, including age, sex and batch effect, were removed by linear regression. We investigated overall cellular compositions of major subtypes, proportional changes of subtypes between case and control, and differential gene expressions and gene co-expression networks in specific subtypes and between case and control.

**Figure 8.**
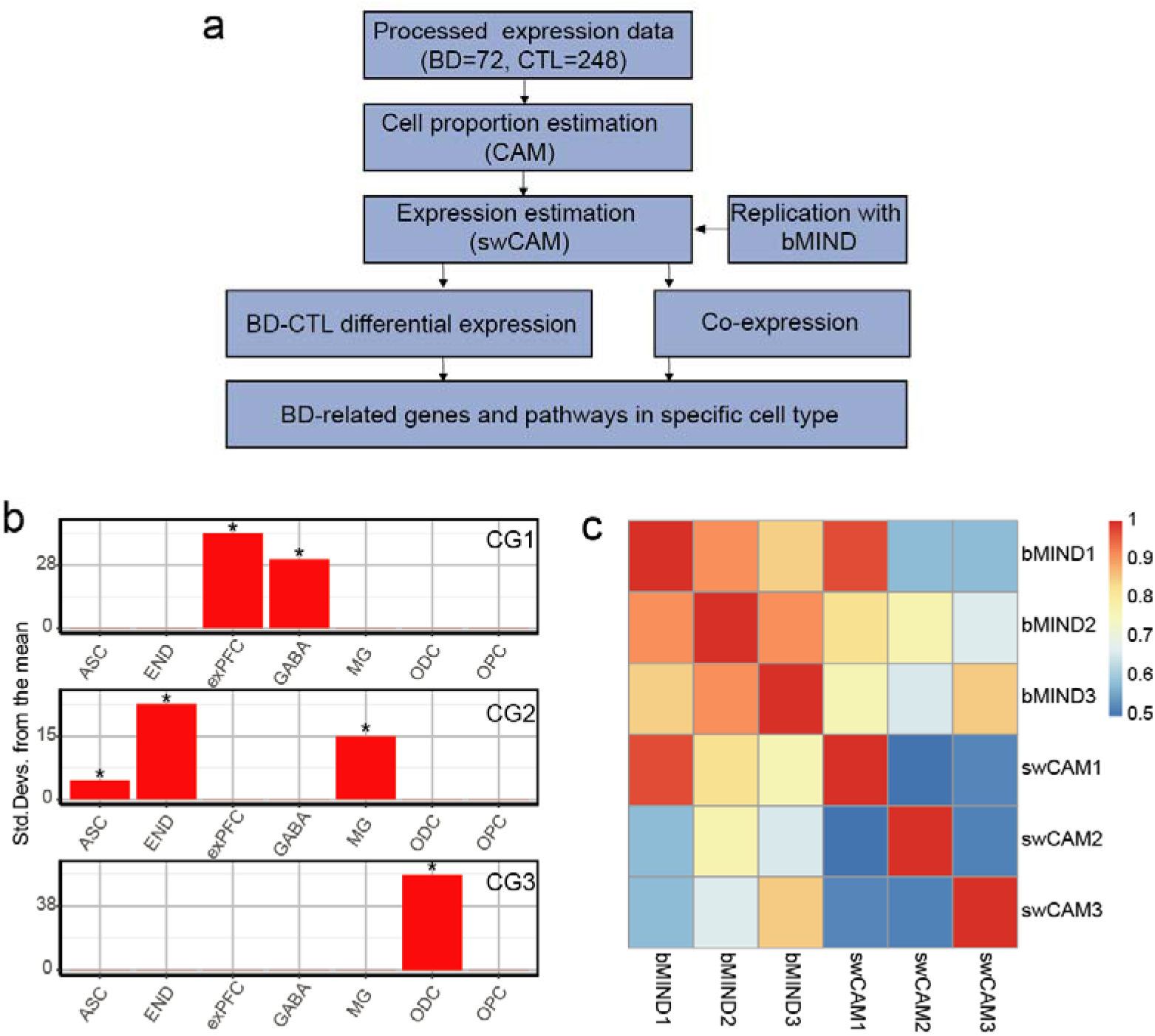
The application of swCAM for biomedical case study. (RNA-seq data of 248 controls and 72 bipolar disorder samples). (a) The analytic pipeline of biological application. (b) The identities of cell groups (CGs) were annotated by EWCE test with snRNA-seq reference from human frontal cortex. The asterisk denotes adjusted p value <0.05. (c) Spearman correlations between CG-specific expressions estimated by swCAM and bMIND.

As shown in Figure 8b, swCAM identified three cell groups (CG) as the subtypes of interest. Using the marker genes of CGs (Figure 8b) and reference of snRNA-seq data from human frontal cortex (Habib, et al., 2017), Expression Weighted Cell Type Enrichment (EWCE) analysis revealed major cellular compositions in CG1 (neuron), CG2 (astrocyte, microglia, and endothelial cell), and CG3 (oligodendrocyte) (Skene and Grant, 2016). We compared the CG-specific expression profiles estimated by swCAM with the replications by bMIND (Wang, et al., 2020). The heatmap of the correlation coefficients over CGs is shown in Figure 8c, with high concordances of 0.96 in CG1, 0.77 in CG2, and 0.85 in CG3, respectively (All Spearman correlation p values <0.05).

Neuron cell group (CG1) showed BD-associated marginal significant changes of cell proportion with a p-value of 0.06 (Figure 9a). The subsequent differential analysis detected 190 down-regulated genes and 21 upregulated genes in CG1 between case and control with FDR <0.05 (Supplementary Table 1 and Figure 9b). Genes related to calcium ion, immune, and mitochondrial systems are downregulated. The further WGCNA analysis identified three differential co-expression modules with FDR <0.05 in CG1 (Supplementary Table 1). The differential expression analysis and WGCNA (e.g. the ME3 and ME19) converged for the downregulation of mitochondria-related genes in neurons, consistent with several previously reported findings about bipolar disorder (Iwamoto, et al., 2005) (Figure 9c-e). Intriguingly, the hub gene of ME3, ATP5H, has previously been reported to be functionally involved in neuron proliferation. Downregulated expression of ATP5H was observed after knockdown of its regulator and resulted in neurogenesis deficit (Su, et al., 2020). The gene functions associated with the differentially expressed or co-expressed genes offer some possible mechanistic insights. For example, the deficiency of mitochondria genes affects the neurogenesis in the early stage, which influence the neuronal functions and contribute to the bipolar disorder pathology.

**Figure 9.**
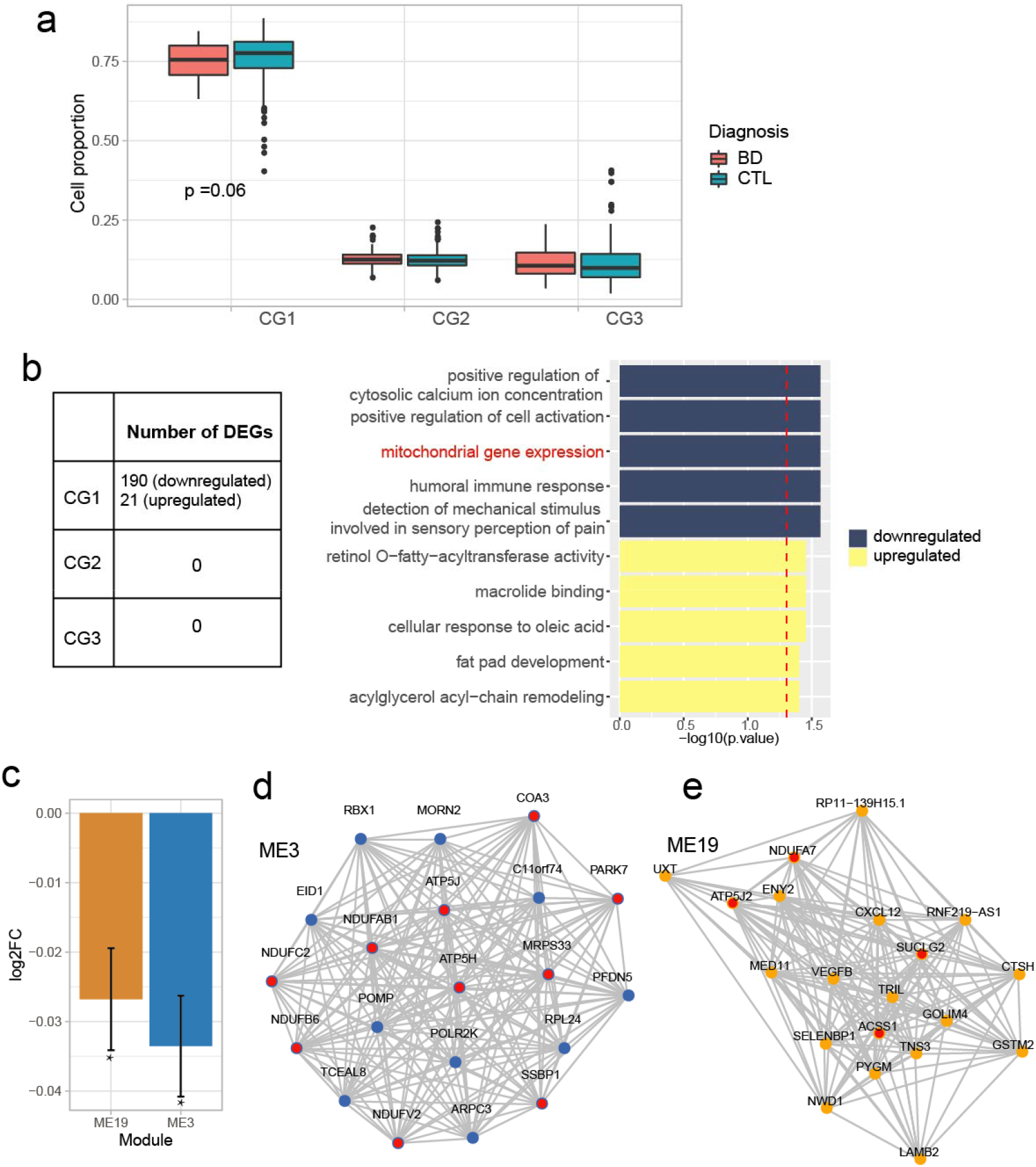
Bipolar disorder associated changes were enriched in the neuron group. (a) The cell proportion changes in each of CGs. P value was from Wilcoxon rank-sum test. (b) CG-specific differential expression results. The threshold for differential expressed genes (DEGs) was FDR corrected p value<0.05. The Red dashed line is the corrected p value of pathway enrichment <0.05. (c) Differential expression of eigengene of two BD-related co-expression modules. The fold change and p value were determined by linear regression. Log2FC= log2 (BD/control). The asterisk denotes corrected p value <0.05. (d-e) Top 20 hub genes in BD-related modules, ranked by module membership (kME). Red nodes are mitochondria-related genes.

### 3.4 Open-source swCAM software tool

We implemented the swCAM workflow in R scripts, an extension of the Bioconductor R package of CAM, freely available at https://github.com/Lululuella/swCAM. A user’s guide and a vignette are provided.

## 4 Discussion

We report a fully unsupervised sample-wise deconvolution method with the R script, swCAM, to estimate unique subtype-specific expressions in individual bulk samples. This method addresses the limitation of population-level deconvolution (averaged across individuals) and enables various subsequent analyses of particular subtypes where higher order statistics within subtypes are required (Zhang and Horvath, 2005; Zhang, et al., 2009). With readily available and tested R scripts, the swCAM tool will allow biologists to perform a deeper characterization of complex tissues in many biomedical contexts. While the principal application here involves gene expression data, the swCAM tool can be readily applied to other molecular omics measurements. The deconvoluted sample-wise transcriptome data enabled many other important downstream analyses, including different expression, co-expression network, and quantitative trait mapping studies.

We emphasize cell groups (multicellular molecularly distinct subtypes) over cell types because the collective and critical effects of cell-cell and cell-matrix interactions are lost when the cellular constituents are studied in isolation. As shown in our real data application, swCAM identified CGs that are composed of one or more canonical cell types. It turns out the brain cells are so diverse that canonical cell types could not reflect the complexity of similarity and dissimilarity among cells. CGs provide a data-driven classification based on transcriptomic signatures that are not obvious by a few cell-type marker genes.

Unsupervised sample-wise deconvolution is mathematically underdetermined (Hastie, et al., 2015). Fundamental to the success of swCAM solution is the nuclear-norm and *ℓ*_2,1_-norm regularized low-rank latent variable modeling. The underlying assumptions in swCAM are biologically plausible because between-sample variations are expectedly governed by a finite number of unique functional modules and each of such modules involves a finite number of genes (Hart, et al., 2015; Zhang and Horvath, 2005). The corresponding subtype expressions estimated by swCAM and bMIND are highly consistent, and the three CGs identified by swCAM are more distinct (lower cross-CG correlation) than that replicated by bMIND (higher cross-CG correlation). It is worth mentioning that experimental results have shown that bMIND provided more accurate sample-level subtype expression estimates as compared to CIBERSORTx (Wang, et al., 2020), where the heuristic algorithm of CIBERSORTx relies on the two assumptions that each gene can be analyzed independently and between-group subtype differential expression is detectable in bulk tissue samples (Newman, et al., 2019).

The cross-validation strategy using random entry exclusion, to determine the proper value of the hyperparameter λ, is adapted from the similar idea in missing value imputation (Hastie, et al., 2015). Our experimental results consistently show a U-curve of the performance index across different values of λ. Note that swCAM is not sensitive to the choice of *λ* within a proper range (plateau of the U-curve). The set-aside masking is a simple yet effective technique for many applications. Here we only demonstrate its application to determine the proper values of *λ* with an experiment on a simulation dataset only to showcase its capability.

There are other ways to provide subtype expressions for individual bulk samples. For example, with population-level subtype expressions obtained using CAM, one-versus-everyone test can be performed to identify a subset of so-called subtype-specific differentially expressed genes (Chen, et al., 2021), and then subsequent analyses of particular subtypes are conducted using the bulk expressions within such a subspace. Alternatively, subtype-abundant bulk samples for each subtype are firstly identified using sample-specific proportions readily provided by CAM, the subtype expression profile for sample *i* and subtype *k* can be approximated by

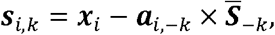

subject to a scaling, where ‘−*k*’ refers to a matrix or vector composed of all other entries except the one associated with subtype *k*. This strategy is consistent with the observation that most methods can extract subtype-specific signals from high abundant subtypes better compared with low abundant subtypes (Rahmani, et al., 2019; Wang, et al., 2020).

