## Supplementary Information for "swCAM: estimation of subtype-specific expressions in individual samples with unsupervised sample-wise deconvolution"

### 1. Introduction

The ability to obtain sample-wise expression variation within each subtype (unique for each individual) is critical prior to infer subtype-specific molecular networks (Junttila and de Sauvage, 2013). Single-cell expression profiling techniques have become popular to investigate cell-type-specific network but may lose critical information of cell-cell interactions and is prone to cell-cycle/state confounders (Buettner, et al., 2015; Gal, et al., 2017). While the existing computational deconvolution methods including our own CAM tool can dissect mixed signals into ‘averaged’ subtype expressions from a population, many subsequent analyses of complex tissues require sample-specific signal deconvolution where each sample is a mixture of ‘individualized’ subtype expression profiles.

In the context of gene regulation and expression, transcriptional regulatory networks connect a finite number of regulatory proteins, such as transcription factors (TFs) and signaling proteins, to the target genes, forming so-called gene co-expression networks or function modules. Based on such mechanistic insights, we exploit the low-rank assumption on the between-sample variations, for each subtype, in our newly proposed swCAM model. While estimating subtype-specific signals from a single mixture is mathematically an underdetermined problem, the low-rank constraints imposed by the swCAM model can aggregate information across genes within function modules to help find a unique and biologically plausible solution.

While general rank minimization is a nonconvex optimization problem (Recht, et al., 2010), minimizing nuclear norm (sum of the singular values over the affine subset) has multiple advantages. Nuclear norm is an accurate convex approximation of the rank function and can be optimized efficiently (Cai, et al., 2010; Candes, et al., 2013; Recht, et al., 2010).

### 2. Methods

Given a penalty parameter $\gamma>0$ (empirically, $\gamma:=1$ generally guarantees good convergence speed), the augmented Lagrangian (ignoring some irrelevant terms) of problem (3) is defined by

$$\mathcal{L}\left( \boldsymbol{U},\boldsymbol{W},\boldsymbol{Z} \right)={\frac{1}{2}\left\| \mathcal{A}\left( \boldsymbol{U} \right)-\boldsymbol{V} \right\|}_{F}^{2}+\lambda\sum_{k=1}^{K} \left\| \boldsymbol{B}_{k}\boldsymbol{W} \right\|_{*}+I_{+}\left( \boldsymbol{B}_{0}\boldsymbol{W} \right)+\frac{\gamma}{2}\left\| \boldsymbol{C}_{1}\boldsymbol{U}+\boldsymbol{C}_{2}\boldsymbol{W}-\boldsymbol{C}_{3}\boldsymbol{-Z} \right\|_{F}^{2}$$

where “$-\gamma\boldsymbol{Z}$”$\in\mathbb{R}^{2KL\times M}$ is the dual variable (or Lagrange multiplier) associated with the constraint $\boldsymbol{C}_{1}\boldsymbol{U}+\boldsymbol{C}_{2}\boldsymbol{W}=\boldsymbol{C}_{3}$. Then, ADMM solves (3) via the following iterative procedure (Chi, et al., 2017):

$$\begin{aligned} \boldsymbol{U}^{q+1}\epsilon\underset{\boldsymbol{U}\in\mathbb{R}^{KL\times M}}{\mathrm{argmin}} \mathcal{L}\left( \boldsymbol{U},\boldsymbol{W}^{q},\boldsymbol{Z}^{q} \right)\#\left( 6a \right) \end{aligned}$$

$$\begin{aligned} \boldsymbol{W}^{q+1}\epsilon\underset{\boldsymbol{W}\in\mathbb{R}^{2KL\times M}}{\mathrm{argmin}} \mathcal{L}\left( \boldsymbol{U}^{q+1},\boldsymbol{W},\boldsymbol{Z}^{q} \right)\#\left( 6b \right) \end{aligned}$$

$$\begin{aligned} \boldsymbol{Z}^{q+1}=\boldsymbol{Z}^{q}-(\boldsymbol{C}_{1}\boldsymbol{U}^{q+1}+\boldsymbol{C}_{2}\boldsymbol{W}^{q+1}-\boldsymbol{C}_{3})\#\left( 6c \right) \end{aligned}$$

where $\boldsymbol{W}^{0}$ can be initialized by $\left[ \boldsymbol{T}_{0}^{T}\boldsymbol{,}\boldsymbol{U}_{0}^{T} \right]^{T}$ with $\boldsymbol{T}_{0}\boldsymbol{=}\boldsymbol{0}_{KL\times M}$ and $\boldsymbol{U}_{0}\boldsymbol{=}\boldsymbol{1}_{M}^{T}\bigotimes\mathrm{vec}\left( {\bar{\boldsymbol{S}}}^{T} \right)$; $\boldsymbol{Z}^{0}$ can be simply initialized by $\boldsymbol{0}_{2KL\times M}$**.** As we will show, both (6a) and (6b) can be solved with closed-form expressions, attributed to the decomposability of ADMM.

Note that (6a) is a column-wise separable optimization problem, so we can decouple w.r.t each column of $\boldsymbol{U}$ (Chi, et al., 2017):

$$\begin{aligned} \boldsymbol{u}_{i}^{q+1}\in\underset{\boldsymbol{u}_{i}\in\mathbb{R}^{KL}}{\mathrm{argmin}} \frac{1}{2}\left\| \boldsymbol{H}_{i}\boldsymbol{u}_{i}\boldsymbol{-}\boldsymbol{v}_{i} \right\|_{2}^{2}+\frac{\gamma}{2}\left\| \boldsymbol{C}_{1}\boldsymbol{u}_{i}+\boldsymbol{y}_{\boldsymbol{i}}^{q} \right\|_{F}^{2}\#\left( 7 \right) \end{aligned}$$

where $\left[ \boldsymbol{y}_{1}^{q},\ldots,\boldsymbol{y}_{M}^{q} \right]\triangleq\boldsymbol{C}_{2}\boldsymbol{W}^{q}-\boldsymbol{C}_{3}\boldsymbol{-}\boldsymbol{Z}^{q}$. The subproblem (7) is an unconstrained quadratic problem, which can be solved by

$$\begin{aligned} \boldsymbol{u}_{i}^{q+1}=\left( \boldsymbol{H}_{i}^{T}\boldsymbol{H}_{i}+\gamma\boldsymbol{C}_{1}^{T}\boldsymbol{C}_{1} \right)^{-1}\left( \boldsymbol{H}_{i}^{T}\boldsymbol{v}_{i}-\gamma\boldsymbol{C}_{1}^{T}\boldsymbol{y}_{\boldsymbol{i}}^{q} \right).\#\left( 8 \right) \end{aligned}$$

The matrix inversion can speed up by

$$\left( \boldsymbol{H}_{i}^{T}\boldsymbol{H}_{i}+\gamma\boldsymbol{C}_{1}^{T}\boldsymbol{C}_{1} \right)^{-1}=\left( \left( \boldsymbol{a}_{i}^{p} \right)^{T}\boldsymbol{a}_{i}^{p}+2\gamma\boldsymbol{I}_{K} \right)^{-1}\bigotimes\boldsymbol{I}_{L}.$$

The right term in (8) can also be simplified as

$$\boldsymbol{H}_{i}^{T}\boldsymbol{v}_{i}-\gamma\boldsymbol{C}_{1}^{T}\boldsymbol{y}_{\boldsymbol{i}}^{q}\boldsymbol{=}\left( \boldsymbol{a}_{i}^{p} \right)^{T}\bigotimes\boldsymbol{x}_{i}^{T}-\gamma\left( \overline{\boldsymbol{y}_{\boldsymbol{i}}^{q}}+\underline{\boldsymbol{y}_{\boldsymbol{i}}^{q}} \right),$$

where $\boldsymbol{y}_{\boldsymbol{i}}^{q}\boldsymbol{=}\left[ \left( \overline{\boldsymbol{y}_{\boldsymbol{i}}^{q}} \right)^{T}\boldsymbol{,}\left( \underline{\boldsymbol{y}_{\boldsymbol{i}}^{q}} \right)^{T} \right]^{T}$with $\overline{\boldsymbol{y}_{\boldsymbol{i}}^{q}}\in\mathbb{R}^{KL}$ and $\underline{\boldsymbol{y}_{\boldsymbol{i}}^{q}}\in\mathbb{R}^{KL}$ being the first and second half vector of $\boldsymbol{y}_{\boldsymbol{i}}^{q}$, respectively (Chi, et al., 2017).

Finally, the column vectors of $\boldsymbol{U}^{q+1}$ in (6a) can be computed fast by

$$\begin{aligned} \boldsymbol{u}_{i}^{q+1}=\mathrm{vec}\left\{ \mathrm{devec}\left\{ \left( \boldsymbol{a}_{i}^{p} \right)^{T}\bigotimes\boldsymbol{x}_{i}^{T}-\gamma\left( \overline{\boldsymbol{y}_{\boldsymbol{i}}^{q}}+\underline{\boldsymbol{y}_{\boldsymbol{i}}^{q}} \right)\left| L,K \right. \right\}\left( \left( \boldsymbol{a}_{i}^{p} \right)^{T}\boldsymbol{a}_{i}^{p}+2\gamma\boldsymbol{I}_{K} \right)^{-1} \right\}\#\left( 9 \right) \end{aligned}$$

To solve (4.6b), we remove some irrelevant terms from its objective function:

$$\begin{aligned} \min_{\boldsymbol{W}\in\mathbb{R}^{2KL\times M}} \lambda\sum_{k=1}^{K} \left\| \boldsymbol{B}_{k}\boldsymbol{W} \right\|_{*}+I_{+}\left( \boldsymbol{B}_{0}\boldsymbol{W} \right)+\frac{\gamma}{2}\left\| \boldsymbol{C}_{1}\boldsymbol{U}^{q+1}+\boldsymbol{C}_{2}\boldsymbol{W}-\boldsymbol{C}_{3}\boldsymbol{-}\boldsymbol{Z}^{q} \right\|_{F}^{2},\#\left( 10 \right) \end{aligned}$$

And then, by defining $\boldsymbol{U}_{k}^{q+1}\in\mathbb{R}^{L\times M},k=1,\ldots,K$ as block matrices from top to bottom in $\boldsymbol{U}^{q+1}\boldsymbol{\in}\mathbb{R}^{KL\times M}$, $\boldsymbol{Z}_{k}\boldsymbol{\in}\mathbb{R}^{L\times M},k=1,\ldots,K$ and $\boldsymbol{Z}_{0}\boldsymbol{\in}\mathbb{R}^{KL\times M}$ as block matrices from top to bottom in $\boldsymbol{Z\in}\mathbb{R}^{2KL\times M}$, respectively (i.e., $\boldsymbol{Z\triangleq}\left[ \boldsymbol{Z}_{1}^{T}\boldsymbol{,\ldots,}\boldsymbol{Z}_{K}^{T}\boldsymbol{,}\boldsymbol{Z}_{0}^{T} \right]^{T}$), we decouple the objective function (10) as functions of $\boldsymbol{T}_{k}\boldsymbol{,}k=1,\ldots,K$ and $\boldsymbol{S}$:

$$\min_{\boldsymbol{W}\in\mathbb{R}^{2KL\times M}} \sum_{k=1}^{K} \left\{ \lambda\left\| \boldsymbol{T}_{k} \right\|_{*}+\frac{\gamma}{2}\left\| \boldsymbol{U}_{k}^{q+1}-\boldsymbol{T}_{k}-\boldsymbol{1}_{M}^{T}\bigotimes{\bar{\boldsymbol{s}}}_{k}\boldsymbol{-}\boldsymbol{Z}_{k}^{q} \right\|_{F}^{2} \right\}+\left\{ I_{+}\left( \boldsymbol{S} \right)+\frac{\gamma}{2}\left\| \boldsymbol{U}^{q+1}\boldsymbol{-S-}\boldsymbol{Z}_{0}^{q} \right\|_{F}^{2} \right\}$$

Therefore, $\boldsymbol{W}^{q+1}$ can be solved by the proximal point algorithm (PPA) (Parikh and Boyd, 2014). Specifically, we have

$$\boldsymbol{W}^{q+1}=\left[ \left( \boldsymbol{T}_{1}^{q+1} \right)^{T},\ldots, \left( \boldsymbol{T}_{K}^{q+1} \right)^{T},\left( \boldsymbol{S}^{q+1} \right)^{T} \right]^{T}$$

in which

$$\begin{aligned} \boldsymbol{T}_{k}^{q+1}\in\underset{\boldsymbol{T}\in\mathbb{R}^{KL\times M}}{\mathrm{argmin}} \lambda\left\| \boldsymbol{T}_{k} \right\|_{*}+\frac{\gamma}{2}\left\| \boldsymbol{U}_{k}^{q+1}-\boldsymbol{T}_{k}-\boldsymbol{1}_{M}^{T}\bigotimes{\bar{\boldsymbol{s}}}_{k}\boldsymbol{-}\boldsymbol{Z}_{k}^{q} \right\|_{F}^{2}\#\left( 11a \right) \end{aligned}$$

$$\begin{aligned} \boldsymbol{S}^{q+1}\in\underset{\boldsymbol{T}\in\mathbb{R}^{KL\times M}}{\mathrm{argmin}} I_{+}\left( \boldsymbol{S} \right)+\frac{\gamma}{2}\left\| \boldsymbol{U}^{q+1}\boldsymbol{-S-}\boldsymbol{Z}_{0}^{q} \right\|_{F}^{2}\#\left( 11b \right) \end{aligned}$$

Note that (4.11a) and (4.11b) are exactly the proximal operators of $\left\| \boldsymbol{T}_{k} \right\|_{*}$ and $I_{+}\left( \boldsymbol{S} \right)$, respectively (Parikh and Boyd, 2014), and their closed-form solutions are given by

$$\begin{aligned} \boldsymbol{T}_{k}^{q+1}=\sum_{\mathcal{l=}1}^{r} \left( \sigma_{k\mathcal{l}}-\frac{\lambda}{\gamma} \right)_{+}\boldsymbol{\mu}_{k\mathcal{l}}\boldsymbol{\nu}_{k\mathcal{l}}^{T},k=1,\ldots,K,\boldsymbol{\#}\left( 12 \right) \end{aligned}$$

$$\begin{aligned} \boldsymbol{S}^{q+1}=\left[ \boldsymbol{U}^{q+1}-\boldsymbol{Z}_{0}^{q} \right]_{+},\#\left( 13 \right) \end{aligned}$$

where the singular value decomposition (SVD) of is performed ahead of the computation of (12), i.e. $\boldsymbol{U}_{k}^{q+1}-\boldsymbol{T}_{k}-\boldsymbol{1}_{M}^{T}\bigotimes{\bar{\boldsymbol{s}}}_{k}\boldsymbol{-}\boldsymbol{Z}_{k}^{q}=\sum_{\mathcal{l=}1}^{r} \sigma_{k\mathcal{l}}\boldsymbol{\mu}_{k\mathcal{l}}\boldsymbol{\nu}_{k\mathcal{l}}^{T}$. A reasonable termination criterion is that the primal residual, $\varepsilon^{pri}=\left\| \boldsymbol{C}_{1}\boldsymbol{U}+\boldsymbol{C}_{2}\boldsymbol{W}-\boldsymbol{C}_{3} \right\|_{2}$, and dual residual, $\varepsilon^{dual}=\left\| \gamma\boldsymbol{C}_{1}^{T}\boldsymbol{C}_{2}\boldsymbol{(}\boldsymbol{W}^{q+1}\boldsymbol{-}\boldsymbol{W}^{q}\boldsymbol{)} \right\|_{2}$, are smaller than a predefined tolerance.

In hyperparameter tuning, (7) is modified to

$$\begin{aligned} \boldsymbol{u}_{i}^{q+1}\in\underset{\boldsymbol{u}_{i}\in\mathbb{R}^{KL}}{\mathrm{argmin}} \frac{1}{2}\left\| P_{\Omega_{i}}^{'}(\boldsymbol{H}_{i}\boldsymbol{u}_{i}\boldsymbol{)-}P_{\Omega_{i}}^{'}(\boldsymbol{v}_{i}\boldsymbol{)} \right\|_{2}^{2}+\frac{\gamma}{2}\left\| \boldsymbol{C}_{1}\boldsymbol{u}_{i}+\boldsymbol{y}_{\boldsymbol{i}}^{q} \right\|_{F}^{2}\#\left( 14 \right) \end{aligned}$$

where $P_{\Omega_{i}}^{'}\left( \cdot\right)=\left[ \boldsymbol{1}_{K}^{T}\bigotimes P_{\Omega_{i}}\left( \cdot\right)^{T} \right]^{T}\in\mathbb{R}^{KL}$ makes all excluded-entry related variables be optimized only by the second term, which is still an unconstrained quadratic problem that can be solved easily. The remaining variables unrelated to excluded entries can still be optimized following (8-9).

Hyperparameter λ actually can be different among different source $k$ and thus the regularization in (2) can be replaced by $\sum_{k=1}^{K} \lambda_{k}\left\| \boldsymbol{T}_{k} \right\|_{*}$ if necessary.

Also, the $\mathcal{l}_{2,1}$ norm of $\boldsymbol{T}$ is defined as

$$\left\| \boldsymbol{T} \right\|_{2,1}\triangleq\sum_{i=1}^{KL} \left\| \boldsymbol{t}_{i} \right\|_{2}$$

accounting for the row-sparsity of $\boldsymbol{T}$. If necessary, the parameter $\delta$ can vary for different rows based on the characteristics of particular genes, such as mean-variance trend. The $\mathcal{l}_{1}$ or $\mathcal{l}_{2}$-norm minimization, as common-used sparsity regularization methods, can also impose entry sparsity constraint on $\boldsymbol{T}$ matrix.

### 3. Software

We implemented the swCAM workflow in R scripts (see Section 5), in addition the Bioconductor R package of CAM (freely available at http://bioconductor.org/packages/debCAM). The R Scripts of the swCAM algorithms are freely available at <https://github.com/Lululuella/swCAM>. A user’s guide and a vignette are provided.

Note that swCAM can be considered an extension of the CAM framework, thus naturally the swCAM pipeline is initialized by the results of CAM (debCAM software package), i.e., the estimates of proportion matrix and subtype-specific expression matrix.

Also note that subsequent analysis of particular subtypes using the results of swCAM is not a formal part of swCAM scripts, e.g., WGCNA (Zhang and Horvath, 2005), DDN (Tian, et al., 2014; Zhang, et al., 2009), or SDEG (Chen, et al., 2021).

### 4. Results

**4.1 Validation on ideal simulation**

Increasing the penalty parameter of the nuclear norm will filter out more noise patterns but at the cost of missing true variation signal. RMSE derived by 10-fold cross-validation strategy is relatively small when λ=1~50 and reach the minimum at λ=5 (Figure S1 and Figure S2). The estimated variation matrices are similar when 1≤λ≤50 (Figure S1e~g), with 12 clear patterns and some artifacts. The artifacts are formed when the signal variation in one subtype spreads to other subtypes for the same genes, which are much lower than detected true signals if λ is not extremely small. The nuclear norm minimization for each subtype’s variation matrix is an effective method to reduce artifacts compared to other regularization schemes

The recovery of sample-specific signals in a subtype is also affected by the mixing proportions of this subtype within the sample. When a subtype accounts for a very small portion in a certain sample, its true signal in this sample will be very weak and thus underestimated (green points in Figure S3). On the contrary, the major subtype in a sample can be estimated very well by swCAM (red points in Figure S3).

**4.2 Assessment on realistic simulation**

RMSE derived by 10-fold cross-validation strategy shows similar trends, see Figure S4. However, the estimated variation matrix by swCAM is blurred by artifacts reflecting noise contamination (Figure S5). Some high-expressed genes have relatively large variance, which could be falsely modeled as subtype-specific signal variations, while the entries with zero value in Ground Truth variation matrix could be overestimated.

We use a $\mathcal{l}_{2,1}$-norm regularization to enforce the sparsity of genes that have signal variation across samples. It is supposed to reduce artifacts while it also follows the assumption that genes contributing to source variation in hidden modules are limited. Figure S6 shows the alleviated artifacts with $\lambda=5$ and $\delta=$10, 1, or 0.1.

Increasing the penalty parameter $\delta$ will force more genes to have zero variance, which suppresses the artifacts and false function modules but brings the risk of missing the true signals. It is critical to propose a parameter tuning method for $\delta$. However, the cross-validation strategy with randomly excluding entries for tuning parameter $\lambda$ is based on the low-rank assumption, where the hidden low-rank patterns can be trained from a part of entries and then used to reconstruct the remaining entries. This strategy is not applicable to $\delta$ selection, which needs further study (Figure S7). The true function modules are correctly detected with $\lambda=5$ and $\delta=1$ or 0.1, where the false module in the first subtype is suppressed when $\delta=1$ (Figure S8). We can still clearly detect 12 functional modules WGCNA constructs weighted networks based on correlation patterns among genes across samples and thus detects function modules of highly-correlated gene sets. The first subtype detects an extra false module, but it is a less significant pattern compared to other modules and can be undetectable with stricter tree height cut threshold.

### 5. Scripts

The script below shows how to obtain some of the results in Figure 6 with proper parameter settings freely available at <https://github.com/Lululuella/swCAM>, supported by the Bioconductor R package of CAM (population-level deconvolution) http://bioconductor.org/packages/debCAM.

```
## Please download ".Rdata" and open "simu.Rproj" for fully replicating Figure 6.

rm(list=ls())
library(gtools)
library(nnls)
library(MASS)
library(truncnorm)
library(ggplot2)
library(scales)
library(gplots)

##########################################
# Simulation data
###########################################

mat <- as.matrix(read.table("GSE19380/plier-mm.matrix.txt",header=T,row.names=1))
Sraw <- mat[,c(1:4, 1:4+8, 1:4+12)]
label <- rep(1:3, each=4)
Smean0 <- sapply(1:3, function(x) rowMeans(Sraw[,label == x]))
L <- n <- 300

set.seed(111)
sampleIdx <- sample(1:nrow(Sraw),n)
Smean <- Smean0[sampleIdx,]

K <- 3
M <- m <- 50 #sample size
r<- 4 # 4 modules in each of 3 types
q<-3
p<- c(0.06,0.06, 0.04,0.04)*n # number of genes involved in 4 modules

SnoiseR <- matrix(rnorm(r*q*m,0,1), r*q, m)
SnoiseR <- SnoiseR - rowMeans(SnoiseR) #center the noise signal
SnoiseL <- matrix(0, K*n, r*q)

ggco <- sample(1:n,q*sum(p)) #co-expressed gene index
A <- rdirichlet(m,c(1,1,1))
start <- 0
#1st cell type
for(i in 1:r){
    coeffi <- runif(p[i],0.15,0.3)
    sign <- rep(c(1,-1), p[i]/2)
    SnoiseL[ggco[start + 1:p[i]], i] <- sign*coeffi
    start <- start + p[i]
}
#2nd cell type
for(i in 1:r){
    coeffi <- runif(p[i],0.15,0.3)
    sign <- rep(c(1,-1), p[i]/2)
    SnoiseL[ggco[start + 1:p[i]] + n, i+r] <- sign*coeffi
    start <- start + p[i]
}
#3rd cell type
for(i in 1:r){
    coeffi <- runif(p[i],0.15,0.3)
    sign <- rep(c(1,-1), p[i]/2)
    SnoiseL[ggco[start + 1:p[i]]  + 2 * n, i+2*r] <- sign*coeffi
    start <- start + p[i]
}

SnoiseVec <-  SnoiseL %*% SnoiseR
Snoise <- t(matrix(c(SnoiseVec),nrow=n))
S.sample <- t(Smean)[rep(1:K,m),] * (1+ Snoise)

X<- c()
for(i in 1:m){
    X <- rbind(X, A[i,,drop=F]%*%S.sample[K*(i-1) + 1:K,])
}
dX <- X - A%*%t(Smean)[1:K,]

Xsd <- runif(m*n, 0.02, 0.05)
Xnoise <- matrix(rnorm(m*n,0,1), m, n)
Xn <- X*(1+Xnoise*Xsd) #end of data simulation

# Assume ture A is known. Otherwize, A needs to be estimated
Sest <- apply(Xn,2, function(x) coef(nnls(A,x)))

################################################################################
# Figure 6a: swCAM with lambda = 5
################################################################################

Sest1 <- Sest
Aest1 <- A

source('sCAMfastNonNeg.R')

rsCAMdtrain <- sCAMfastNonNeg(Xn, Aest1, Sest1, iter = 1,  r = 1, lambda = 5,
                              iteradmm=10000, silent = T, eps = 1e-10)
breaks = seq(-1, 1, length.out=129)

ggcoReorder <- ggco
ggcoReorder <- c(ggcoReorder, (1:n)[-ggco])

optvar0 <- Smean[,rep(1:K, each = M)]

optvar <- cbind(rsCAMdtrain$W[1:n,],rsCAMdtrain$W[n+1:n,],rsCAMdtrain$W[2*n+1:n,])
heatmap.2((t((optvar/optvar0)[c(ggcoReorder),])), trace="none",keysize = 1,
        Rowv = F,Colv = F,#col=redblue(64),
        add.expr = abline(v=c(0+0.5, L+0.5), h= c(0+0.5, M+0.5, 2*M +0.5, 3*M+0.5),lwd=1,col='black'),
        labRow = FALSE, labCol = FALSE,margins = c(2, 2),
        col=bluered(128), key=F, breaks=breaks, symkey=FALSE, density.info="none",
        lmat=rbind(4:3,2:1), lhei=c(0.1,4), lwid=c(0.1,4))


################################################################################
# Figure 6b: swCAM with lambda = 5, delta = 1
################################################################################

source('sCAMfastNonNeg_NN_L21.R')

rsCAMdtrain <- sCAMfastNonNeg_NN_L21(Xn, Aest1, Sest1, iter = 1,  r = 1, lambda = 5, delta = 1,
                                     iteradmm=10000, silent = T, eps = 1e-10)
optvar0 <- Smean[,rep(1:K, each = M)]

optvar <- cbind(rsCAMdtrain$W[1:n,], rsCAMdtrain$W[n+1:n,], rsCAMdtrain$W[2*n+1:n,])
heatmap.2((t((optvar/optvar0)[c(ggcoReorder),])), trace="none",keysize = 1,
        Rowv = F,Colv = F,#col=redblue(64),
        add.expr = abline(v=c(0+0.5, L+0.5), h= c(0+0.5, M+0.5, 2*M +0.5, 3*M+0.5),lwd=1,col='black'),
        labRow = FALSE, labCol = FALSE,margins = c(2, 2),
        col=bluered(128), key=F, breaks=breaks, symkey=FALSE, density.info="none",
        lmat=rbind(4:3,2:1), lhei=c(0.1,4), lwid=c(0.1,4))

################################################################################
# Figure 6c: 10-fold cross-validation to determine lambda
################################################################################

Kfold <- 10
set.seed(111)

nall <- ncol(Xn) * nrow(Xn)
tmpseq <- rep(1:Kfold, nall %/% Kfold)
if (nall %% Kfold > 0) {
    tmpseq <- c(tmpseq, 1:(nall %% Kfold))
}
sample_exp <- matrix(sample(tmpseq), nrow(Xn))

source('sCAMfastNonNegNA.R')

Sest1 <- Sest
Aest1 <- A

library(doSNOW)
cl <- makeCluster(Kfold, type = "SOCK")
registerDoSNOW(cl)

lambdaAll <- c(10000, 5000, 2000, 1500, 1000, 800, 500, 400, 300, 200, 150, 100, 80, 50, 40, 30, 20, 10 , 8, 5 , 2, 1, 0.5, 0.2, 0.1, 0.05, 0.01, 0.001, 0.0001, 0.00001)

ptm <- proc.time()
res <- foreach (ifold = 1:Kfold, .combine=rbind) %dopar% {
    warmstart <- matrix(0, L * K, M)
    Xtrain <- Xn
    Xtrain[sample_exp == ifold] <- NA
    errlambda <- c()

    for (lambda in lambdaAll) {
        rsCAMdtrain <- sCAMfastNonNegNA(Xtrain, Aest1, Sest1, iter = 1,  r = 1, lambda = lambda,
                                        iteradmm=10000, silent = T, eps = 1e-10, warm.start=warmstart)

        warmstart <- rsCAMdtrain$W[1:(L*K),]

        Xest <- c()
        for(i in 1:M){
            tmps1 <- Sest1 + t(matrix(rsCAMdtrain$W[1:(K*L),i], nrow = ncol(Xn)))
            tmps2 <- t(matrix(rsCAMdtrain$W[1:(K*L) + K*L,i], nrow = ncol(Xn)))
            tmps <- (tmps1 + tmps2) /2
            Xest <- rbind(Xest, Aest1[i,,drop=F]%*%tmps)
        }
        errlambda <- c(errlambda, sum((Xest[sample_exp == ifold] - Xn[sample_exp == ifold])^2),
                       rsCAMdtrain$epPri, rsCAMdtrain$epDual, rsCAMdtrain$iterrun)
    }
    errlambda
}
proc.time() - ptm

stopCluster(cl)

rmse <- res[,seq(1,ncol(res),4)]
rmse <- sqrt(rmse/ (M  * L /Kfold))

self_fun <- function(x) {sqrt(sum(x^2 *(M  * L /Kfold) ) / (M*L))}

df.cv <- data.frame(x=factor(rep(lambdaAll, each=nrow(rmse))),y=c(rmse))
ggplot(subset(df.cv, x %in% c(10000, 5000, 2000, 1500, 1000, 800, 500, 400, 300,
                              200, 150, 100, 80, 50, 20, 10 , 8, 5 , 2, 1, 0.5,
                              0.2, 0.1, 0.05, 0.01, 0.001, 0.0001, 0.00001)),
       aes(x=x, y=y)) + geom_boxplot() + theme_bw(base_size = 16) +
       stat_summary(fun.y=self_fun, geom="line", aes(group=1),  colour="blue") +
       theme(axis.text.x = element_text(angle = 90, hjust = 1, vjust = 0.5, size=10))+
       xlab("lambda") + ylab("RMSE")
```

### 6. Supplementary Table 1

See separate excel spreadsheets.

### 7. Supplementary Figures

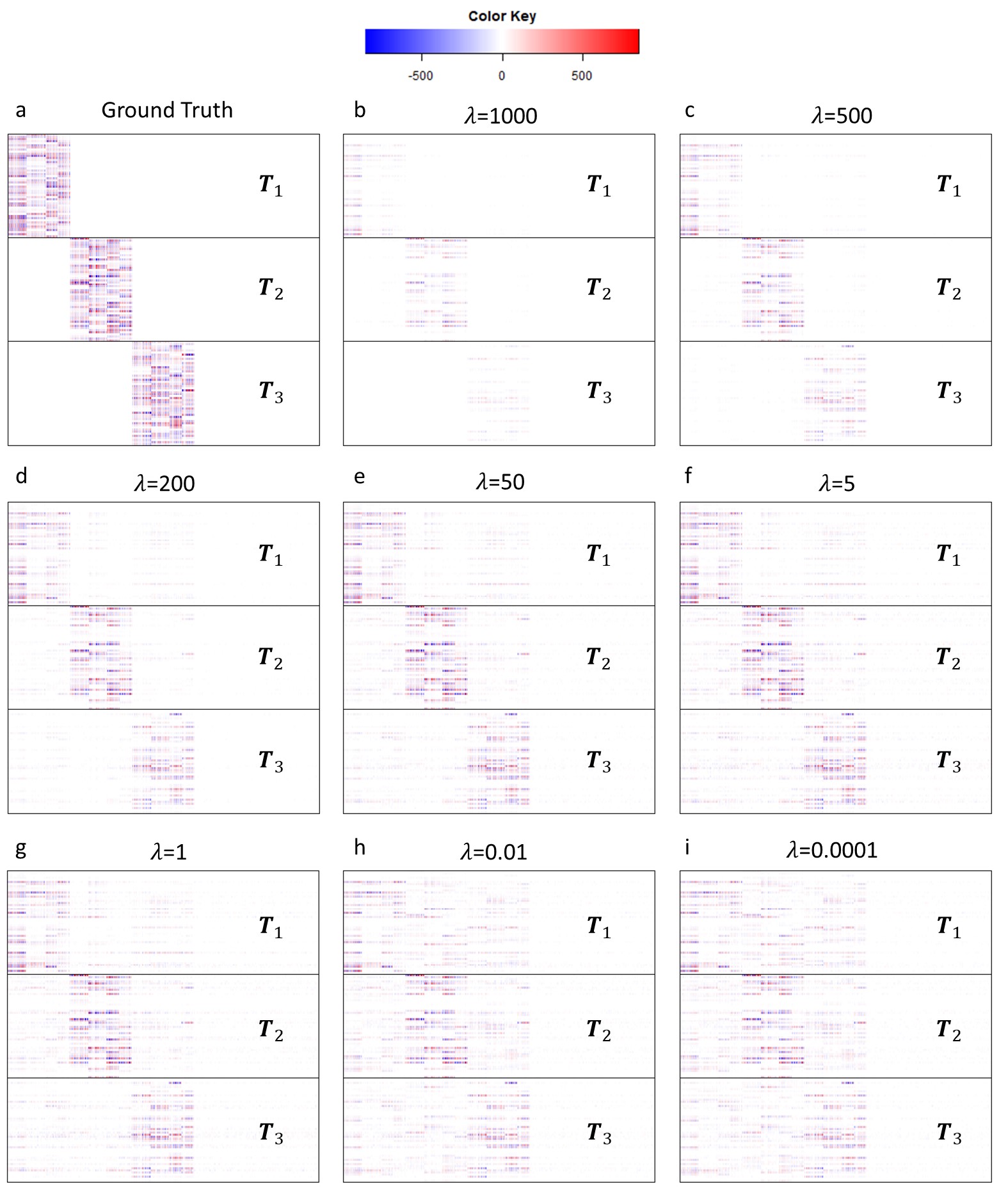

**Figure S1.** Heatmap of estimated $T$ matrix with varied $\lambda$ parameters compared to ground truth in the ideal simulation. Increasing the penalty parameter of the nuclear norm will filter more noise but at the cost of the possibility of missing true signal variation.

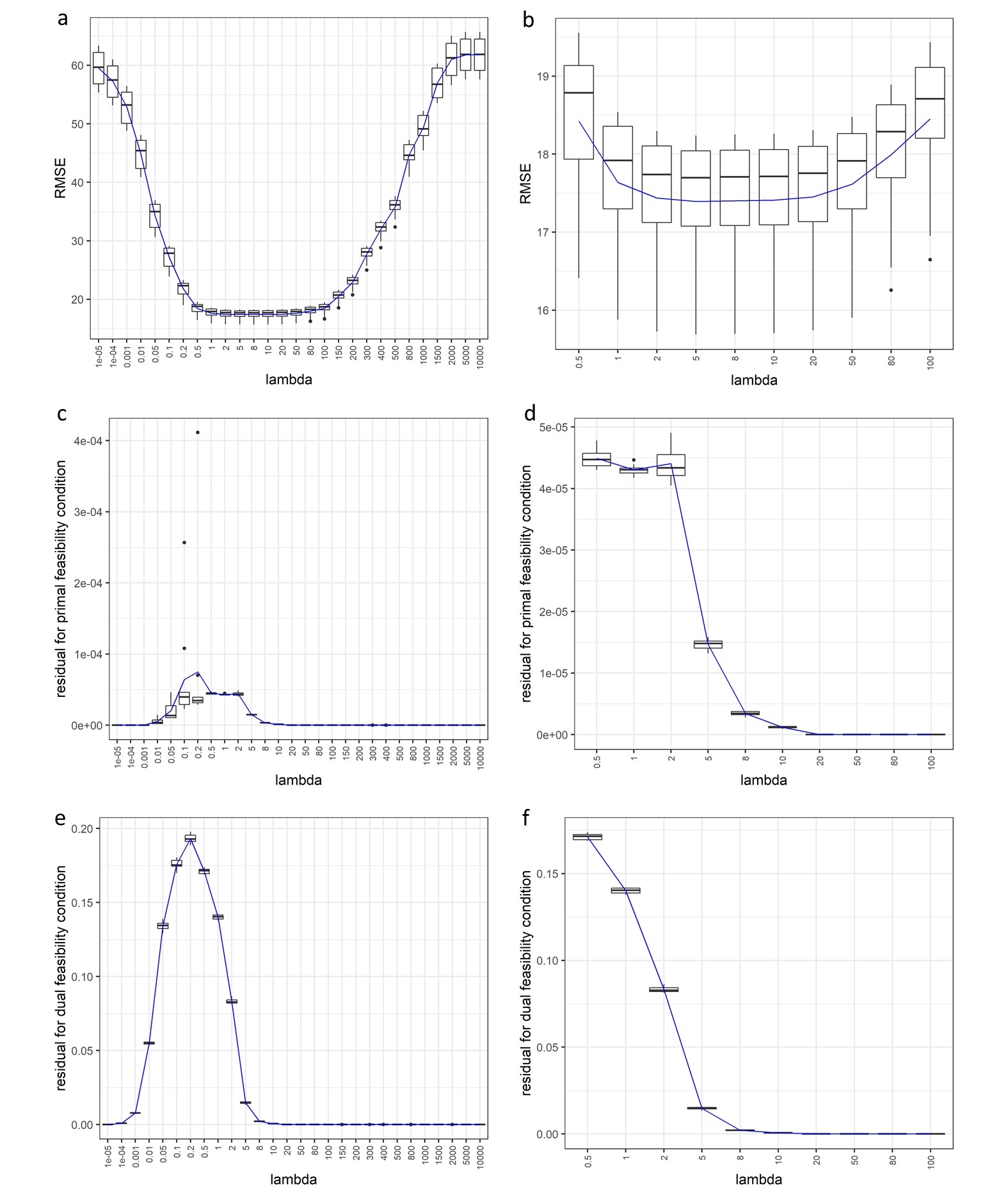

**Figure S2.** 10-fold cross-validation results under different $\lambda$ parameter in the ideal simulation. (a) RMSE; (c) Residuals for primal feasibility condition; (e) Residuals for dual feasibility condition; (b), (d), (f) are zoomed curves of (a), (c), (e).

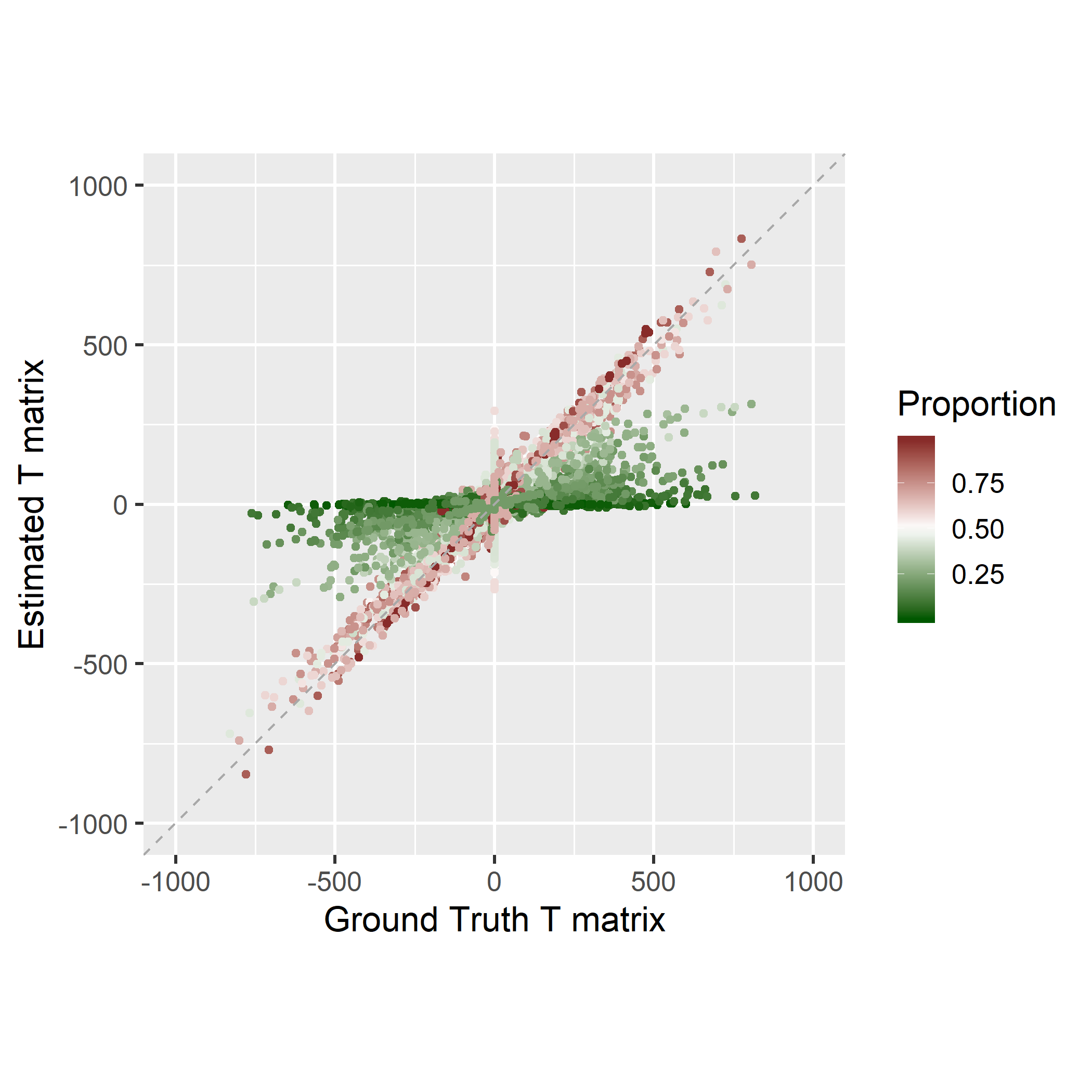

**Figure S3.** Estimated $\boldsymbol{T}$ matrix versus ground truth when $\lambda$=5 in the ideal simulation. The mixing proportions associated with estimated entries are colored to show the sample-specific expression estimations for high-proportion subtypes can be estimated more accurately than those for low-proportion ones.

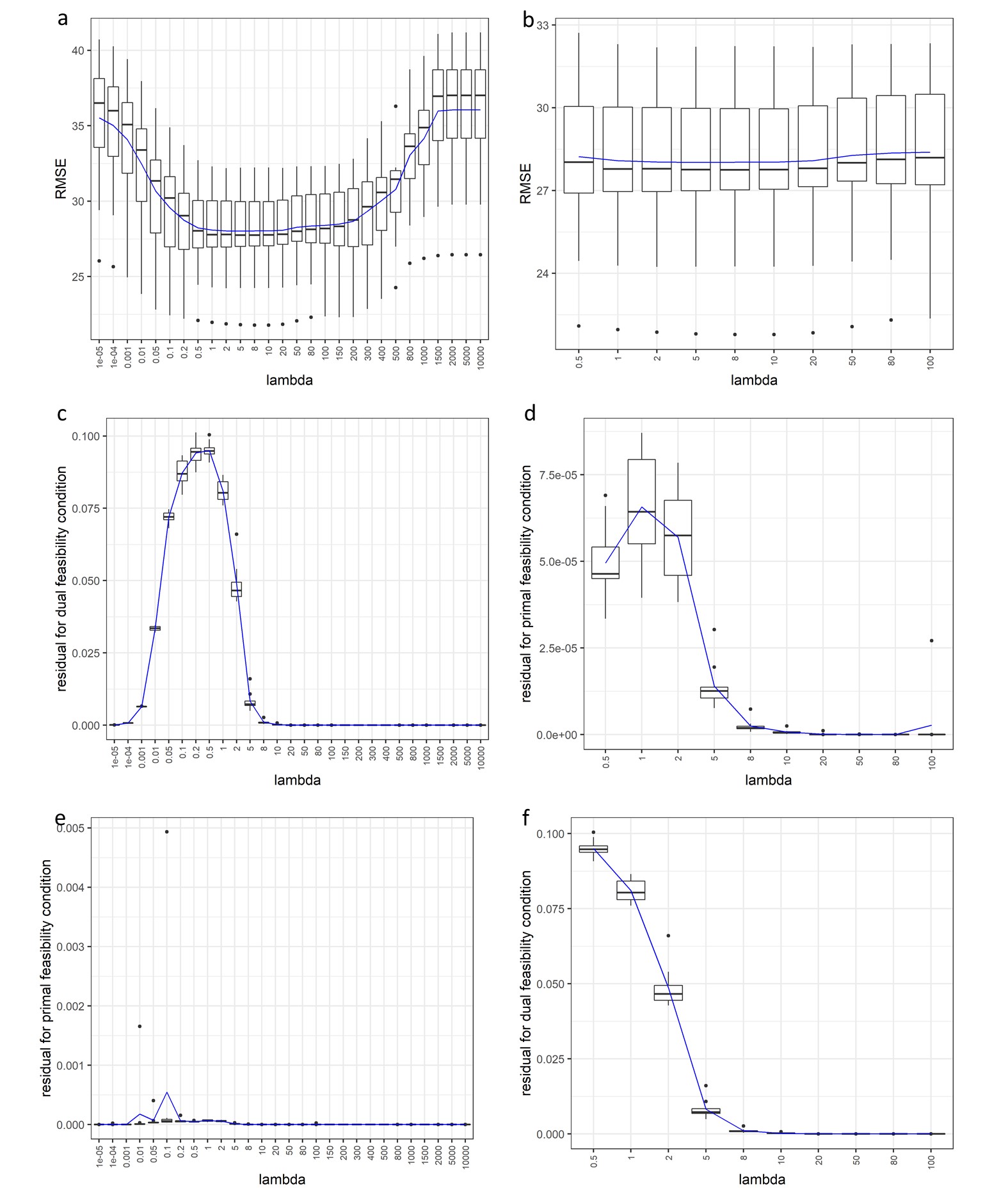

**Figure S4**. 10-fold cross-validation results under different $\lambda$ parameter in the realistic simulation. (a) RMSE; (c) Residuals for primal feasibility condition; (e) Residuals for dual feasibility condition; (b), (d), (f) are zoomed curves of (a), (c), (e).

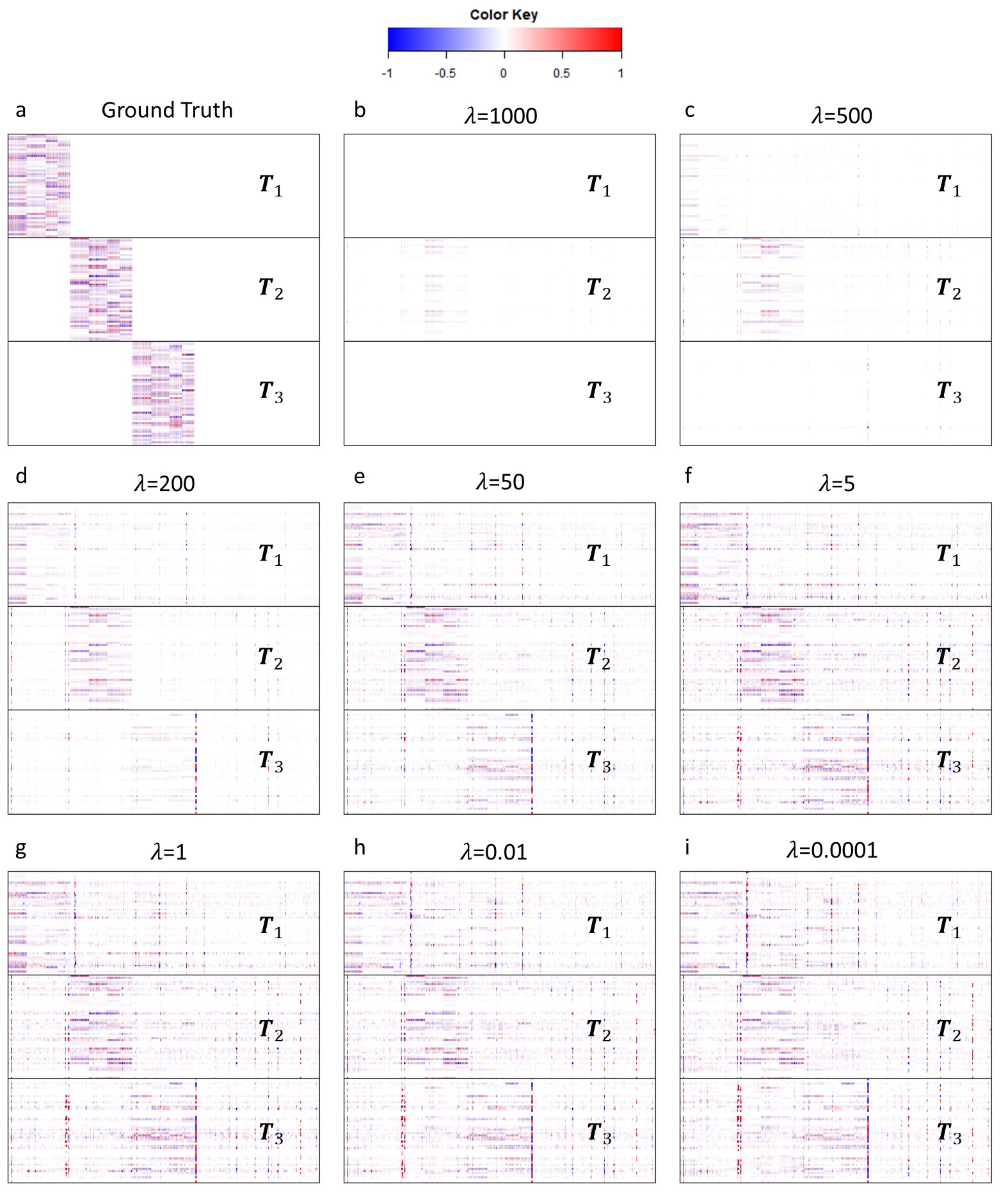

**Figure S5.** Heatmap of estimated $\boldsymbol{T}$ matrix scaled by associated means compared to ground truth in the realistic simulation with varied $\lambda$ parameters. Increasing the penalty parameter of the nuclear norm will filter more noise but at the cost of the possibility of missing true variation signal.

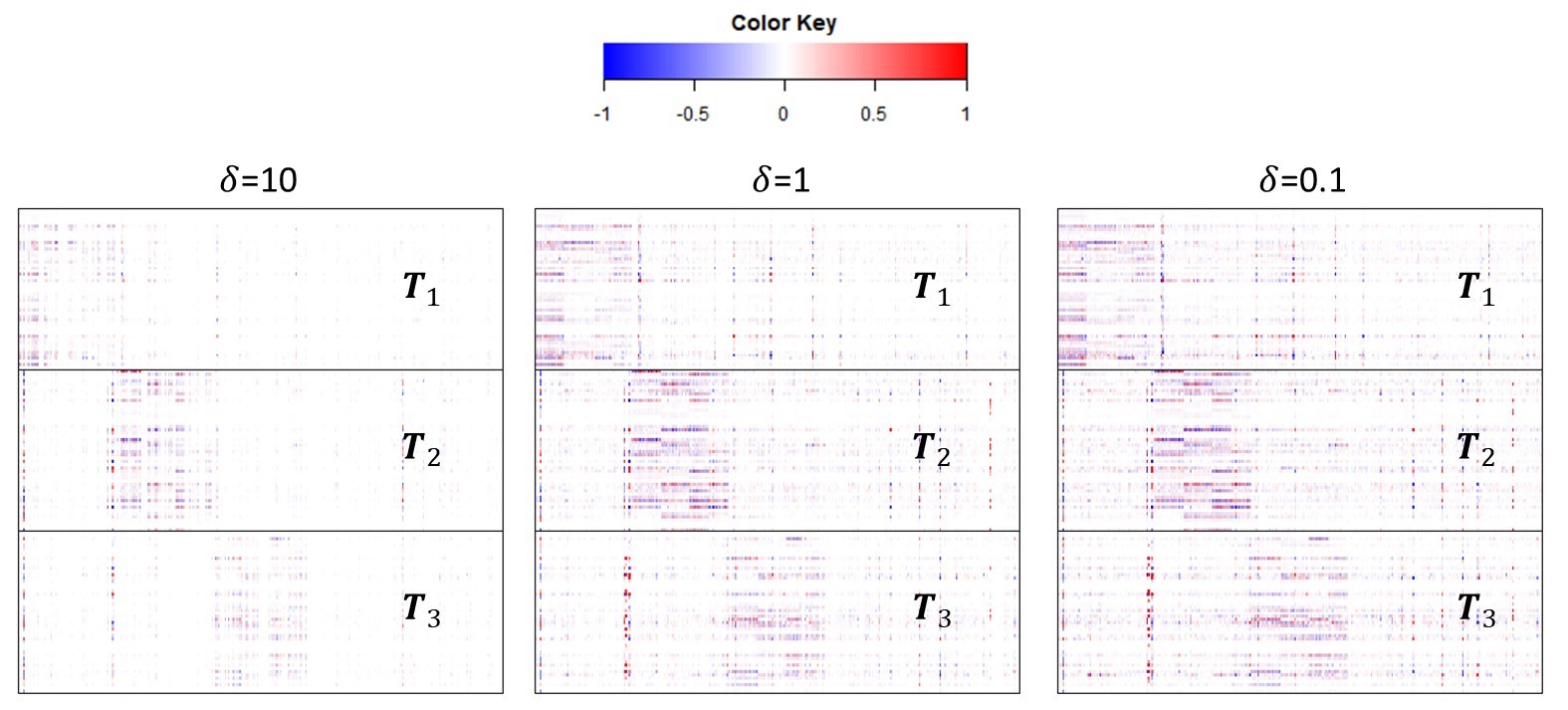

**Figure S6.** Heatmap of estimated T matrix scaled by associated means compared to ground truth in the realistic simulation with $\lambda=5$ and varied $\delta$. Increasing the penalty of L_2,1_ norm will enforce more zero columns in $\Delta S_{k}$ matrix.

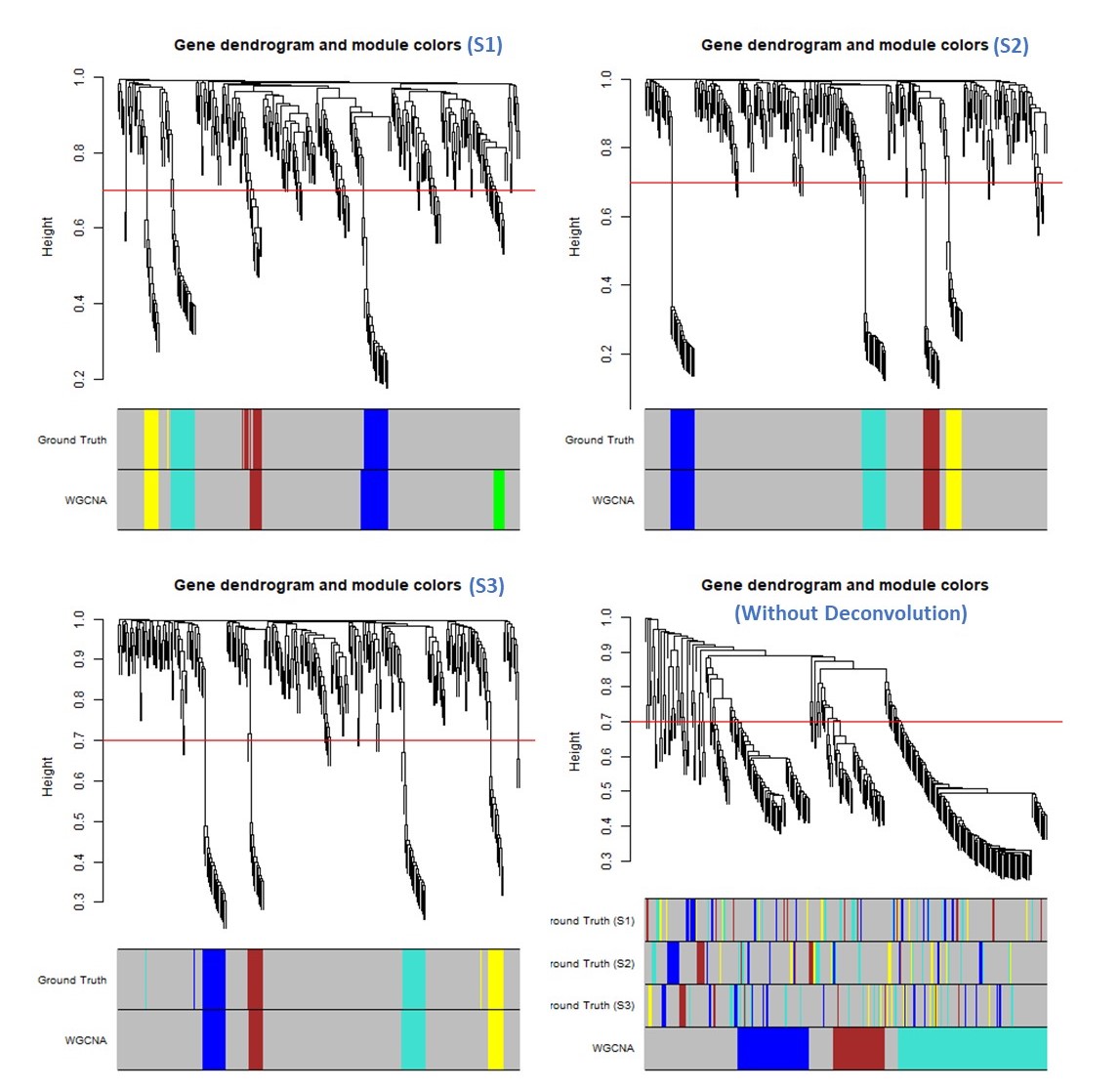

**Figure S7.** Gene co-expressed function modules detected by WGCNA on swCAM estimated sample-specific expression for each subtype (a~c) or on originally observed expressions without deconvoluton (d). (Network interconnectedness is measured by topological overlap; cutHeight = 0.7; minSize = 8).

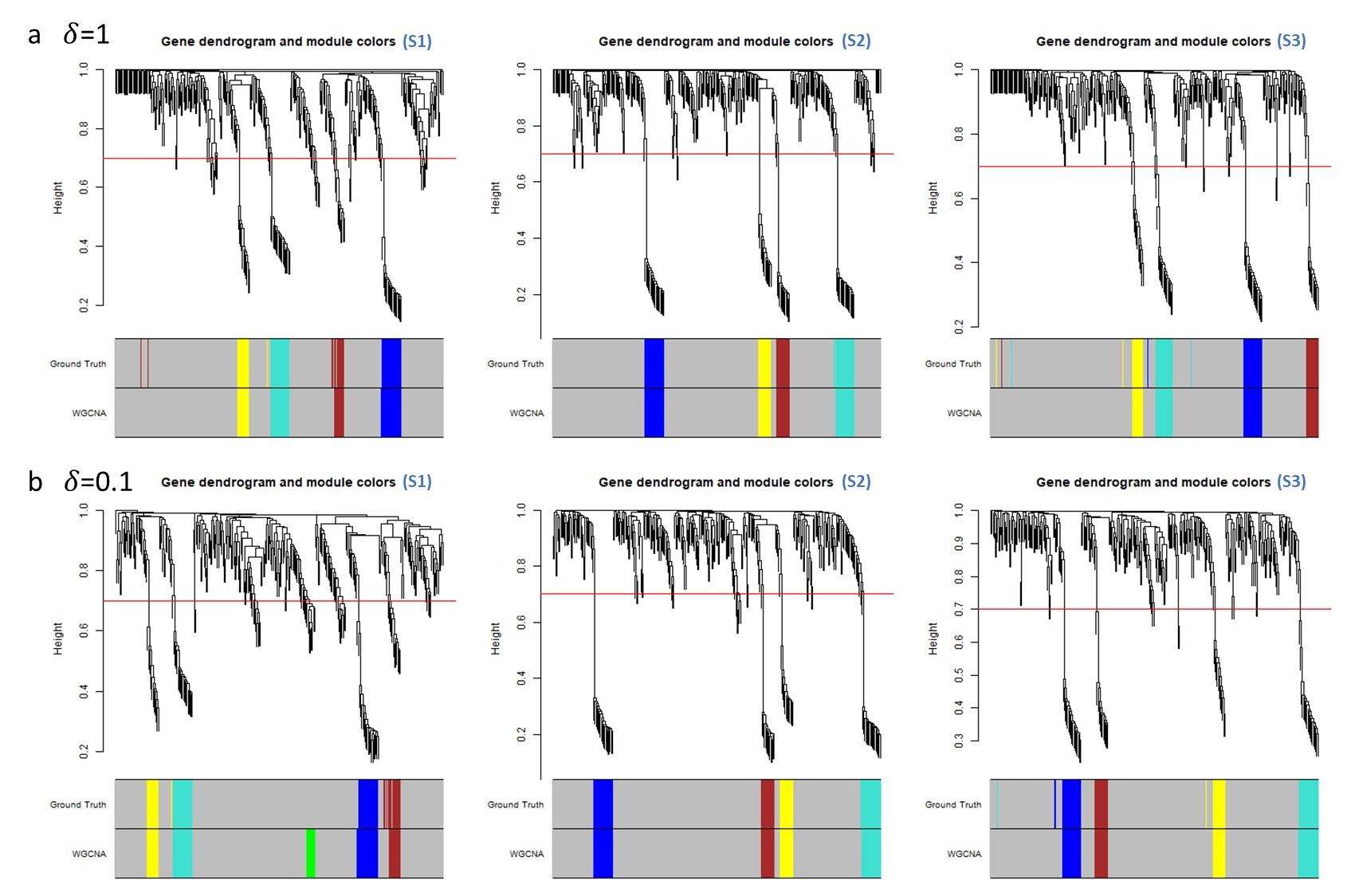

**Figure S8.** Gene co-expressed function modules detected by WGCNA on swCAM estimated sample-specific expression for each subtype with λ=5 and δ=1 or 0.1. (Network interconnectedness is measured by topological overlap; cutHeight = 0.7; minSize = 8).
